# The landscape of tertiary lymphoid structures in endometrial cancer revealed through harmonized multi-level transcriptomics

**DOI:** 10.1101/2025.08.20.671212

**Authors:** Marta Requesens, Koen Brummel, Arkajyoti Bhattacharya, Feija Somefun, Nienke van Rooij, Annechien Plat, Dimas M. X. van der Hall, Annegé Vledder, René Wardenaar, Diana C.J. Spierings, Eugene Berezikov, Tjalling Bosse, Floris Foijer, Joost Bart, Hans W. Nijman, Rudolf S.N. Fehrmann, Marco de Bruyn

**Affiliations:** Department of Obstetrics and Gynaecology, University Medical Center Groningen, University of Groningen, Groningen, The Netherlands; Department of Medical Oncology, University Medical Center Groningen, University of Groningen, Groningen, The Netherlands; European Research Institute for the Biology of Ageing, University Medical Center Groningen, University of Groningen, Groningen, The Netherlands; Department of Pathology, Leiden University Medical Centre, Leiden, The Netherlands; Department of Pathology and medical biology, University Medical Center Groningen, University of Groningen, Groningen, The Netherlands

**Keywords:** Tertiary lymphoid structures, anti-tumor immunity, spatial transcriptomics, Independent-component analysis, immune-checkpoint blockade

## Abstract

Tertiary lymphoid structures (TLSs) are ectopic lymphoid tissues that form within tumors. TLSs are associated with improved prognosis and better responses to immune checkpoint inhibitors (ICI) in cancer. However, the contribution of genetic, immune, and (micro)environmental factors to TLS formation and function remain incompletely understood.

Using harmonized bulk and single-cell RNA-sequencing coupled with spatial transcriptomics, we present a comprehensive characterization of tumor-associated TLSs in endometrial cancer (EC). Our integrated datasets reveal the cellular architecture of TLSs, their organizing principles—including chemokine-receptor gradients—and TLS activity compared to paired tumor-draining lymph nodes, as well as spatial preferences in IgA/IgG plasma cell and T cell distribution.

We identified a consensus set of transcriptional components that independently predict the presence of TLS in EC. Furthermore, we demonstrate that clinical neoadjuvant ICI in EC patients shifts the cellular balance in favor of immunogenic *CXCL13*+ *PDCD1*+ T cells, providing a mechanistic insight into ICI-induced TLSs formation. Our data offers a systems-level understanding of TLSs and a roadmap for their therapeutic exploitation.

## INTRODUCTION

Tertiary Lymphoid Structures (TLSs) of various complexities arise in peripheral tissues in response to persistent immunological stimuli or chronic inflammation, such as autoimmunity^1^, organ transplant^2^, and cancer^3^. TLSs have been identified in a wide range of cancers and at different stages of disease, yet they display considerable heterogeneity in both location and morphology. Based on their cellular composition and degree of organization, the maturation status of TLSs ranges from small diffuse clusters to highly organized lymphoid and stromal structures, referred to as mature TLS (mTLS). mTLSs exhibit an architecture similar to that of secondary lymphoid organs, characterized by an inner CD20+ B cell follicle surrounded by a distinct CD3+ T cell zone^3^. Additionally, mTLSs are defined by the presence of CD21+ follicular dendritic cells (FDCs) and CD4+ T follicular helper cells (Tfh), which contribute to the formation of germinal center (GC) reactions. Within these reactions, B cells undergo clonal selection, somatic hypermutation, class switch recombination, and differentiation into antibody-secreting plasma blasts^4^. Furthermore, mTLSs are suggested to support T cell development, promote CD8+ T cell effector phenotypes^5^, and shape antigen-specific adaptive immune responses *in situ* through LAMP3+ dendritic cells (DCs) – T cell interactions^6^, ultimately promoting superior overall anti-tumor immunity^3^.

With few exceptions, the presence of TLSs correlates with clinical benefit across cancer types in both primary and metastatic disease^7^, including but not limited to melanoma^8^, lung cancer^9–11^, hepatocellular carcinoma^12^, pancreatic cancer^13^, colorectal cancer^14,15^, ovarian cancer^5,16^, and endometrial cancer (EC)^17,18^. In fact, mTLSs predict responses to ICI in multiple cancer types^8,19–23^ and have been shown to be a superior predictive factor compared to T cell infiltration^8^ or PD-L1 status^22^. Despite these strong clinical associations with patient outcomes, the underlying mechanisms by which TLSs contribute to anti-tumor immunity are still poorly understood. A better understanding of how TLSs shape *in situ* immunity and drive ICI outcomes could pave the way for TLS-inducing therapies and improve clinical ICI outcomes.

We previously demonstrated that the presence of ≥1 TLS within or around the tumor in EC patients is independently associated with improved survival across the four different EC molecular subtypes^17^. This makes EC a valuable model for gaining deeper insights into TLS dynamics in anti-tumor immunity. In this study, we applied spatial transcriptomics (10x Genomics Visium) and single-cell RNA-sequencing (scRNAseq) to characterize the landscape of TLSs in EC. We utilized paired tumor-draining lymph nodes (TDLN) to establish the relative transcriptional and proliferative profiles of TLS and the corresponding TDLN-GC. We integrated these spatial and scRNAseq datasets with bulk gene expression profiles from The Cancer Genome Atlas (TCGA) to identify statistically independent transcriptional patterns of TLSs in EC. Finally, we propose a potential mechanistic basis for ICI-mediated TLS induction by analyzing how neoadjuvant ICI in EC patients influences the activities of these transcriptional patterns. Our findings highlight the organizing principles of TLS and suggest potential new targets for the therapeutic exploitation of TLSs in immunotherapy.

## RESULTS

We performed spatial transcriptomics (ST) on six EC cases with histologically proven mTLS (Figure 1A and S1). These included one ultra-mutated POLE-mutant tumor (POLEmut), two mismatch repair-deficient (MMRd) tumors, two TP53-mutant (p53mut) tumors and one tumor of the no specific molecular profile (NSMP) subtype (a KRAS-G12V mutant). In addition, we leveraged scRNAseq data from eight endometrial tumors in the human cell atlas (HCA)^24^, complemented with in-house generated scRNAseq of tumor-infiltrating lymphocytes (TILs) from an additional four EC patients (Figure 1A and Figure S2A). We specifically sequenced CD3+, CD19+, and CD3-CD19-CD45+ cells separately to enrich for rarer immune subtypes, such as mature dendritic cells (mDC *LAMP3*; Figure S2B). Within the ST sections, the expression of canonical TLS markers (*CD19, BCL6, MKI67*, and *CR2*) correlated with pathologist annotation of TLS on the H&E sections (Figures 1B-C and S1A-B), as did global cell type inference using the scRNAseq reference (Figure 1D and S1C).

**Figure 1.**
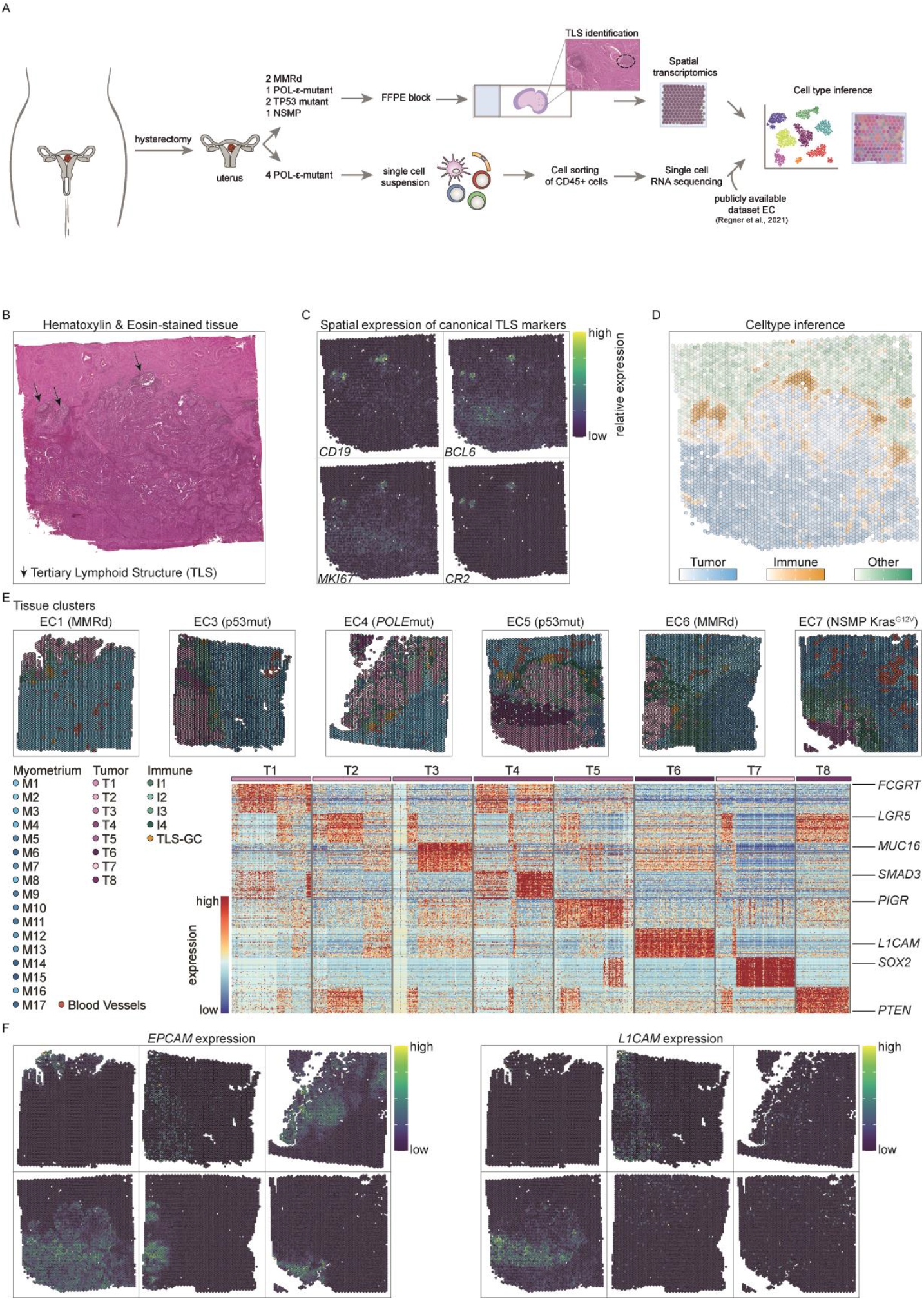
Spatial Transcriptomics of EC sections containing TLSs. (A) Schematic overview of the work-flow. ST or scRNAseq was performed on EC samples and integrated with a publicly available dataset to infer the cell types. (B) Representative H&E staining of an EC section in which ST was performed. Black arrows indicate the pathologist’s annotation of TLS. (C) Spatial gene expression of *CD19, BCL6, MKI67* and *CR2* and (D) celltype inference depicting tumor (blue), immune (orange) and other (green) cell type ST spots across a representative EC section. (E) Clusters of ST spots depicted across samples EC1-EC7. Heatmap of differential gene expression across the 8 tumor clusters. (F) *EPCAM* (left) and *L1CAM* (right) spatial gene expression across the 6 EC samples. ST: Spatial Transcriptomics; H&E: hemotoxylin and eosin; EC: endometrial cancer; TLS: tertiary lymphoid structures; MMRd: mismatch-repair deficient; NSMP: no specific molecular profile; EPCAM: epithelial cell adhesion molecule; L1CAM: L1 cell adhesion molecule.

Hierarchical clustering of ST spots resulted in 18 myometrial clusters (including blood vessels), eight tumor clusters and five immune clusters (including a TLS-GC cluster) (Figure 1E). The epithelial tumor marker *EPCAM* was expressed in all tumor clusters across sections (Figure 1F), while the well-established prognostically unfavorable biomarker *L1CAM* was differentially expressed (DE) in tumor cluster T6 (Figure 1E). This cluster was prevalent in the p53mut tumors and exhibited heterogeneous expression (Figure 1F). Gene set enrichment analysis (GSEA), using a guilt-by-association (GBA) approach, revealed additional heterogeneity among the tumor clusters, primarily related to epithelial differentiation, angiogenesis, cell cycle, and immune-related biological processes (Figure S3). DE genes from the myometrial clusters revealed subtle spatial differences in extracellular matrix molecules, endothelial markers, and immune cell infiltration (Figure S4A). The immune clusters were largely defined by their immune cell composition, although cluster I4 was also substantially enriched for genes related to collagen formation and deposition (Figure S4B). The TLS-GC cluster exhibited a distinct gene expression profile characterized by high expression of genes essential for TLS formation (*CXCL13, LTB*), FDC markers (*CR2*), and B cell-related function (*CD19, MS4A1, PAX5, IKAROS, CD79, CD72*), indicating a robust presence of key components involved in the development and maintenance of TLSs and active B cell processes (Figure S4).

Thus, our dataset accurately captures TLS and EC biology and serves as a novel resource for studying TLS in particular, and EC in general.

### Spatial distribution of B cell and Ig-receptors with TLS

A subsequent granular cell type inference (Figure 2A and S5) and canonical marker analysis (Figure S6) confirmed that GC-B cells (GCBs) and naïve/memory B cells (NMBCs) predominantly populated the TLS-GC clusters, whereas antibody-secreting cells (ASCs) were more evenly distributed across clusters (Figure 2A, and S5). In line with ongoing GC activity and class switch recombination, we observed higher expression of *IGHD* and *IGHM*, mostly found on naïve B-cells^25^, in close proximity to the TLS-GC cluster, with a sharp declined outside the TLS-GC border (Figure 2B). By contrast, *IGHG1, IGHG2* and *IGHG4* genes remained largely constant up to 1mm outside of TLS-GCs, while *IGHA1* expression increased up to 1mm (Figure 2B), consistent with ASCs being present in various cell clusters within the tumor microenvironment (TME) (Figures 2A). Immunohistochemistry (IHC) validation using a sequential section of EC3 confirmed that IgA accumulated more distally from TLS, whereas IgG was more evenly distributed (Figure 2C). Notably, analysis of individual TLS-GC clusters revealed a degree of isotype heterogeneity, even within a single patient (Figure S7).

**Figure 2.**
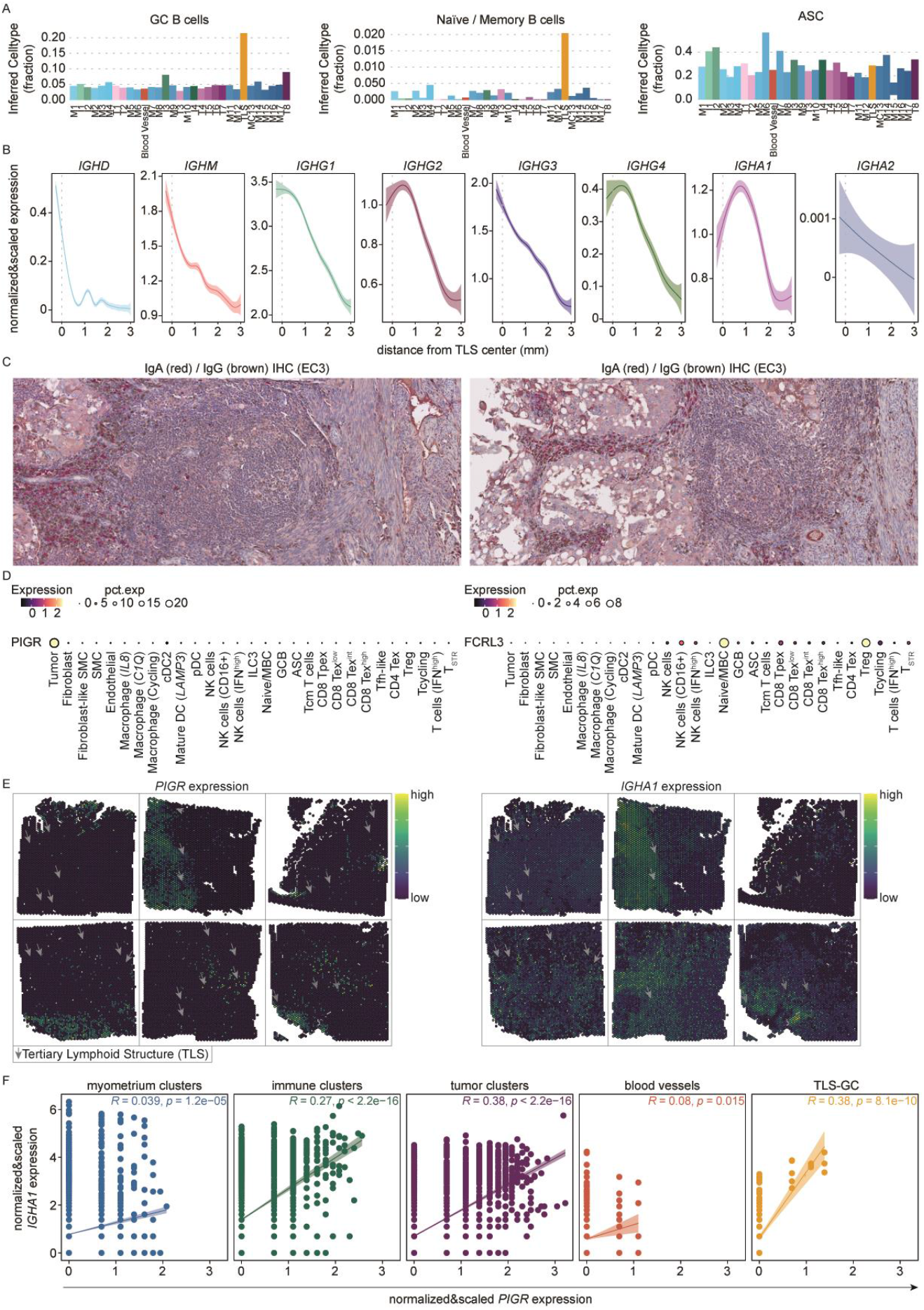
B cell immunity associated with TLSs. (A) Inferred fraction of GCBs, naïve/MBs and ASCs across clusters. (B) Distance of B-cell derived Ig gene expression in relationship to the TLS center (mm). (C) Representative image of double IHC staining of IgA (red) and IgG (brown) of a TLS in EC3. (D) Expression of *PIGR* and *FCRL3* across cell types identified. (E) Spatial gene expression of *PIGR* and *IGHA1* across all EC sections. Grey arrows indicate the location of the TLSs. (F) Correlation plots of the normalized and scaled expression of *IGHA1* and *PIGR* along different clusters found in the EC-samples. GCB: germinal center B-cell; naïve/MB: naïve/ memory B-cell; ASC: antibody-secreting cell; Ig: immunoglobulin; IHC: immunohistochemistry; T_STR_: stressed T cells; T reg: regulatory T cells; T_ex_: exhausted T cells, T_fh_-like: follicular helper T cells, T_pex_: precursor-exhausted T cells; T_cm_: central-memory T cells; ILC3: innate-lymphoid cells group 3; NK: natural killer cells; pDC: plasmacytoid dendritic cells; SMC: smooth muscle cells.

Recent studies have shown that IgA responses are capable of targeting intracellular (neo)antigens^26^, promoting T cell immunity through the polymeric immunoglobulin receptor (PIGR)^27^, and converting regulatory T (Treg) cells into an inflammatory phenotype via binding to FCRL3 on the Treg cell surface^28^. Accordingly, we observed *PIGR* expression in EC tumor cell clusters in our scRNAseq data, and *FCRL3* expression in Treg cells and NMBCs (Figure 2D). Spatially, *PIGR* was expressed in three of the six tumors and co-localized with *IGHA1*-expressing B cells (Figure 2E and 2F). By contrast, and consistent with the immune cell expression observed in the scRNAseq data, *FCRL3* was expressed in ST sections in and around TLS (Figure S8A), which we confirmed through immunofluorescence analysis (Figure S8C). Notably, *FCRL3* expression did not spatially correlate with *IGHA1* (Figure S8A and B).

We analyzed the migratory cues that might underly these differences in the spatial orientation of *IgA*+ or *IgG*+ ASCs. The only previously published ST study of TLSs found that fibroblast-produced CXCL12 was likely to be a key factor in organizing ASC migration from TLSs to tumors^29^. We therefore analyzed expression of CXCL12 and other chemokines in spatial relationship to TLSs. Consistent with earlier findings^30^, the uterine myometrium was a significant source of *CXCL12* within our spatial dataset (Figure S9A). Nevertheless, we also observed *CXCL12* expression in pathologist-annotated tumor-associated stromal regions, as well as in the fibroblast population in the scRNAseq dataset (Figure S9B). ASCs in our dataset did not express other previously reported migratory ASC receptors (Figure S10).

It therefore remains to be determined whether CXCL12-CXCR4 is a pan-cancer migratory pathway that organizes the spatial location of TLS-derived ASC, or whether additional cancer-specific migratory cues exist.

### Spatial architecture of TLS reveals sites of T cell help and chemokine-dependent organization

Expanding outward from the TLS center, we observed that cluster immune 3 (I3) was uniquely distributed around TLS-GC clusters (Figures 3A and S11A). Cluster I3 was primarily comprised of exhausted (*PD-1*+, *CTLA-4*+ and *LAG-3*+) *CXCL13*+ CD4 T and Treg cells (Figure 3B and S5), consistent with the role of CXCL13+ cells in promoting B-cell recruitment and the formation of a TLS T cell zone^31^. Cluster I3 was also characterized by a relatively high fraction of mature LAMP3+ DCs (Figure S5), and it was the only cluster where CD4+ T cells and DCs were spatially correlated (P=0.011). LAMP3+ DCs were marked by canonical CD maturation markers, including *CD86, CD83* and *CCR7* (Figure 3C). In line with recent reports that the maturation of DCs to a CCR7+ LAMP3+ state requires interaction with CD4+ T cells, we observed significant enrichment of the DC “help” signature (Figure 3D) in these cells^32^. “Helped” DCs were further characterized by increased expression of immune checkpoint molecules *CD274* and *CD200*, recently described to have a pivotal role in ICI response (Figure 3D)^33^. Consistent with cluster I3, which is rich in CD4-mDC, serving as a potential niche for T cell priming or reactivation, the Gini-TCR index—a measure of TCR clonality diversity—was higher in regions proximal to TLSs (Figures 3E and S11B). Notably, this higher diversity was not a direct reflection of increased CD4+ or CD8+ T-cell densities (Figures S11C-D). While CCR7+ LAMP3+ DCs are intrinsically migratory towards TDLN, recent studies have identified a subset of these cells retained in the TME^33^. We speculated that TLS might “divert” these cells from the TDLN, leading us to analyze intra-tumoral chemokine expression gradients. The *CCL19*/*CCL21-CCR7* and the *CCL22-CCR4* ligand-receptor pairs showed a spatial gradient with respect to TLS proximity (Figures 3F and S12). *CCL19* and *CCL22* were produced by CCR7+ LAMP3+ mDC (Figure S10), with CCR4 prominently expressed on exhausted *CXCL13*+ CD4 T and Treg cells (Figure 3G), suggesting the formation of self-organizing antigen-presenting cell (APC) niches near TLSs.

**Figure 3.**
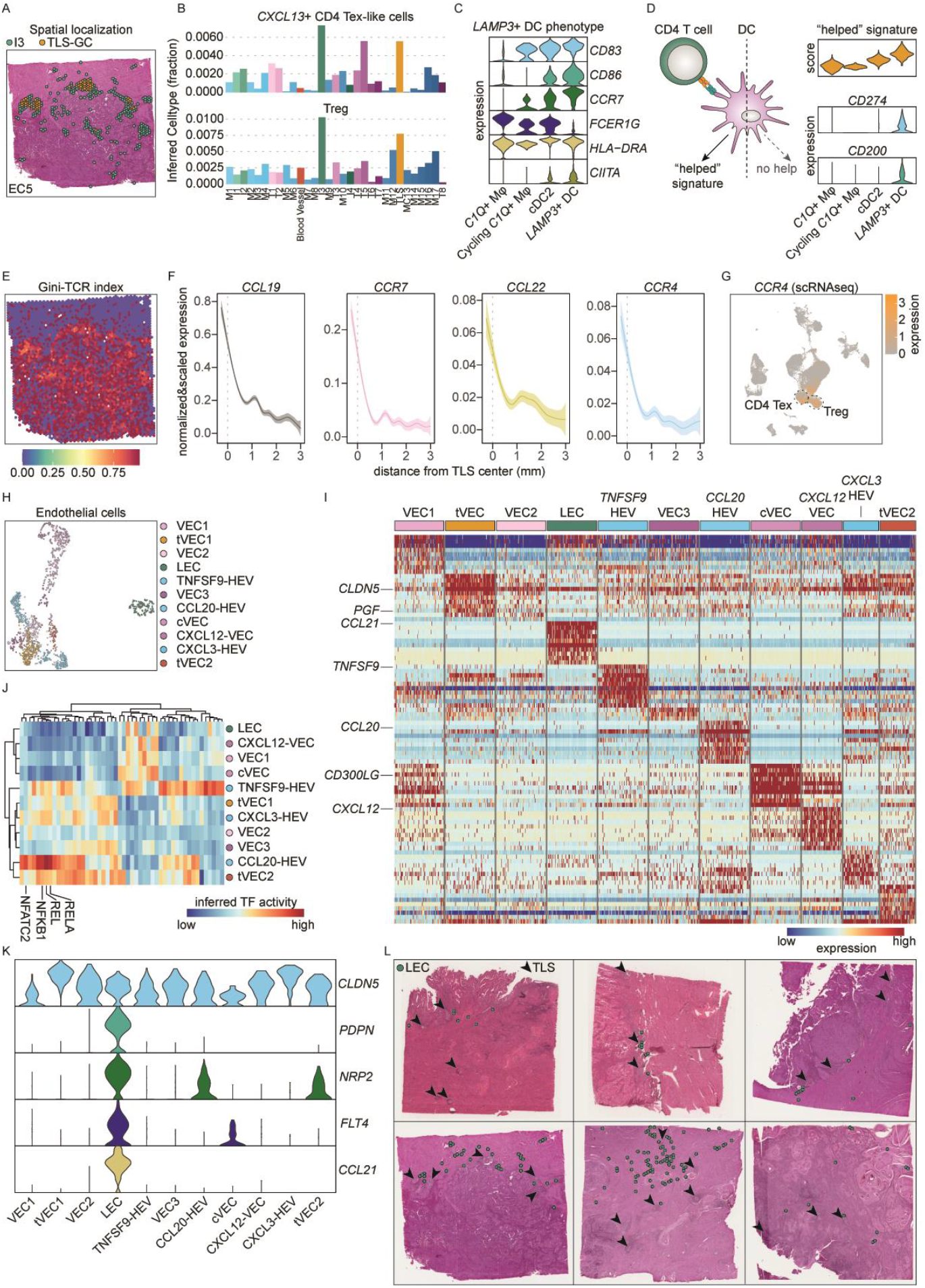
Characterization of TLS-associated cell subsets. (A) Representative example of spatial colocalization of CD4-containing cluster I3 and the TLS cluster. (B) Inferred fraction of *CXCL13+* CD4+ T exhausted cells and regulatory T cells across clusters. (C) Gene expression of canonical DC maturation markers across different subsets of macrophages and DCs. (D) Score of the “helped signature” and expression of CD200 and CD274 across different subsets of macrophages and DCs. (E) Gini-TCR index in a representative EC-slide. (F) Expression of *CCL19, CCR7, CCL22*, and *CCR4* in relationship to the distance from the TLS center (mm). (G) UMAP projection of CCR4 expression across the different cell populations. (H) UMAP depicting subclusters of endothelial cells. (I) Heatmap showing the differential gene expression of endothelial cell clusters depicted in (H). (J) Heatmap showing an inferred substantial activity of NF-κB transcription factors in HEC clusters. (K) Violin plots illustrating canonical paracortical LEC gene expression in EC LECs. (L) Spatial localization of the single cell LEC cluster in EC-samples. TLS: tertiary lymphoid structure; T_ex_: T exhausted; DC: dendritic cell; TCR: T-cell receptor; scRNAseq: single-cell RNA-sequencing; HEC: high-endothelial cell; LEC: lymphatic endothelial cell; TF: transcription factor.

In contrast to CCL19, we observed *CCL21* expression not in mDC but in a subset of endothelial cells (Figure S10). In lymph nodes, CCL21+ lymphatic endothelial cells (LECs) are spatially colocalized with high-endothelial cells (HECs) in the (para-)cortex and are thought to mediate rapid lymphocyte egress and ingress, thus facilitating antigen scanning by naïve and memory lymphocytes^34,35^. To investigate whether a similar organization exists in TLSs, we analyzed endothelial cells in more detail. We identified seven vascular endothelial (VEC) subclusters characterized by expression of canonical markers *PECAM*1+ *PROX1-*, a capillary VEC (cVEC) cluster identified by *CD300LG+ NOX5*+^36,37^, two PGF+ tip VEC (tVEC) clusters involved in vessel sprouting, a single *PECAM1+ PROX1+* lymphatic endothelial cell (LEC) cluster^36^ and three high endothelial venule (HEV) subclusters (Figure 3H). Consistent with earlier studies, we noted no *CCL21* mRNA expression in human *ACKR1+ SELE+* HEVs ^36,38^ (Figure 3I). Differential gene expression analysis revealed several immune-modulating HEV states, including *TNFRSF9*+ (41BBL), *CCL20*+, and *CXCL3*+ HEVs, as well as a cluster of (*ACKR1-SELE-*) *CXCL12*+ VEC cells (Figure 3I and 3J). In line with the proposed trans-differentiation of VEC to HEV through progressive TNF/LTβR-induced NFκB-signaling^39^, we inferred substantial activity of NF-κB-related transcription factors (TFs) in HEV clusters (Figure 3J). In addition to expression tVEC marker PGF, tVEC cluster 2 was associated with REL, RELA, NFKB1, and NFATC2 HEV-TF activity, possibly indicating ongoing HEV remodeling in the TME^40^ (Figure 3J). LECs were further characterized by expression of *PDPN, NRP2, FLT4*, and *CCL21* (Figure 3K), consistent with the cortical and paracortical LEC phenotype in lymph nodes, as well as markers previously reported for capillary LECs^35,41^. Spatial analysis of the LEC cluster revealed a distribution consistent with a capillary phenotype, with sites throughout the myometrium, around blood vessels, and colocalized near TLS (Figure 3L).

Together, our data provide a potential mechanism for the – previously observed^17^ – preferential formation of TLS in the myometrium of EC patients and their self-organization during maturation. Whether similar principles underly preferential intra- or peritumoral localization of TLS remains to be determined.

### Higher proliferative and germinal center activity in TLS-GCs compared to TDLN-GCs

The contribution of TLS-GC activity versus TDLN-GC activity to tumor immunity remains largely unknown. Therefore, we performed ST analysis on several paired and one unpaired TDLNs from patients who underwent clinically indicated lymphadenectomy (n=3). As a control for lymph node germinal center activity, we included tonsil tissue. Similar to what was observed in the TLS-GC cluster, the expression of canonical GC markers *CD19* and *CR2* correlated with pathologist annotation of the H&E sections (Figures 4A-B and S13A-D). However, we noted a striking absence of GC activity markers *BCL6* and *MKI67* in TDLN GCs (Figures 4B and S13B-C), implying reduced activity and proliferation in TDLN GCs. This observation was further supported by substantially lower G2M and S scores in the TDLNs compared to the control tonsil (Figure 4C). As expected, hierarchical clustering of ST spots revealed a high prevalence of GC clusters in the tonsil (Figures 4D and S13E). By contrast, we observed substantial fibrosis and the ubiquitous presence of macrophages in the TDLN (Figures 4D and S13E). Regarding the retention of mDC in the TME due to *CCL19/21* gradients, we did not observe obvious defects in *CCL19/21* production in the TDLNs, as both HEV and DC clusters expressed *CCL19* and *CCL21* (Figure S13F).

**Figure 4.**
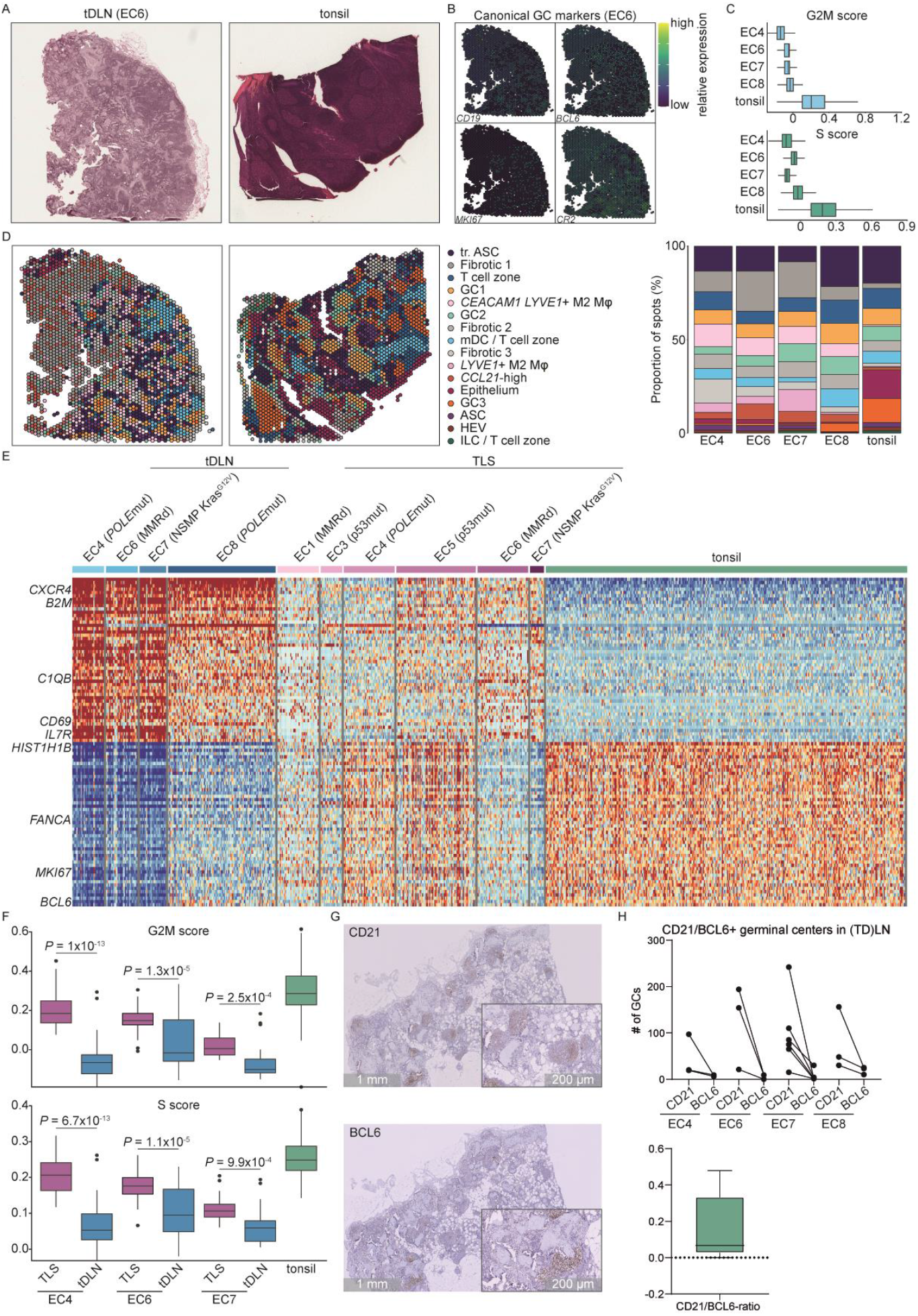
Germinal center activity and proliferation in TLS and TDLN. (A) Representative H&E-section of a TDLN from EC8 and a tonsil used for reference. (B) Spatial gene expression of *CD19, BCL6, MKI67*, and *CR2* in the TDLN from EC8. (C) G2M- and S-scores EC TDLN and tonsil as reference. (D) Clusters of ST spots depicted across TDLN (EC8) and tonsil and the % of spots in every cluster across the different samples. (E) Heatmap of differential gene expression for indicated proliferation-associated genes for TDLN, TLS and tonsil samples. (F) Box-and-whisker plots depicting G2M- and S-scores in TLS-GC, TDLN-GC and tonsil-GC. (G) Representative image of IHC of TDLN germinal centers with CD21 and BCL6. (H) Paired dot plot depicting the presence of CD21+ and BCL6+ GCs across TDLN in EC patients; Box-and-whisker plot showing the BCL6/CD21 ratio in GCs of TDLN. H&E: hematoxylin and eosin; TDLN: tumor-draining lymph node; ST: Spatial transcriptomics; IHC: immunohistochemistry; GC: germinal center.

Differential gene expression analysis of TDLN clusters revealed that GC cluster 3, the most prevalent GC cluster in tonsils, was the only GC cluster characterized by genes associated with cell proliferation (Figure S14). TDLN were enriched for non-proliferative GC clusters 1 and 2. To determine the composition and activity of TLS-GCs compared to TDLN and tonsil GCs, we compared all four lymph node GC clusters to TLS-GC clusters. TLS-GCs exhibited a gene expression profile intermediate to the tonsil and TDLN GCs, with more frequent expression of *MKI67, BCL6, FANCA*, and *HIST1H1B* compared to TDLN GCs (Figure 4E). Moreover, paired analysis revealed that G2M and S scores were significantly higher in TLS-GCs compared to paired TDLN-GCs (Figure 4F). We confirmed these observations using IHC for CD21 and BCL6 on the TDLNs used for ST, as well as on all other lymph nodes from these patients, and found a lower number of BCL6/CD21+/+ GCs relative to CD21+ GCs in TDLN-GCs (Figure 4G-H).

All in all, these data support ongoing and higher GC activity and function in TLS-GCs when compared to paired TDLNs.

### Transcriptional patterns of the TME associated with TLS

Having established the composition and relative transcriptional functionality of TLSs, we next sought to determine transcriptional patterns consistently associated with TLSs in EC. We developed a strategy to overcome the inherent limitations of small sample sizes in whole-slide spatial transcriptomics. In brief, we applied consensus-independent component analysis (c-ICA) to gene expression profiles from the TCGA, which contains expression levels of 20,392 genes across 10,817 samples. C-ICA decomposes gene expression profiles into additive independent components, referred to as transcriptional components (TCs), with each TC being as statistically independent as possible from the others. We then determined which TCs displayed an association with TLSs by analyzing a subset of 218 EC patients from TCGA, for whom TLS status had been previously annotated by a pathologist review^17^. For a subset of 39 TCs the activity was associated with the presence of TLSs (referred to as TLS-associated TCs; false discovery rate 5%, confidence level 80% in permutation-based multiple testing framework; Source Data; Figure 5A). Of these, 22 TCs exhibited significantly higher activity in samples with TLSs compared to those without (Figure 5B). The remaining 17 TCs captured transcriptional effects that were more active in samples lacking TLSs (Figure 5B).

**Figure 5.**
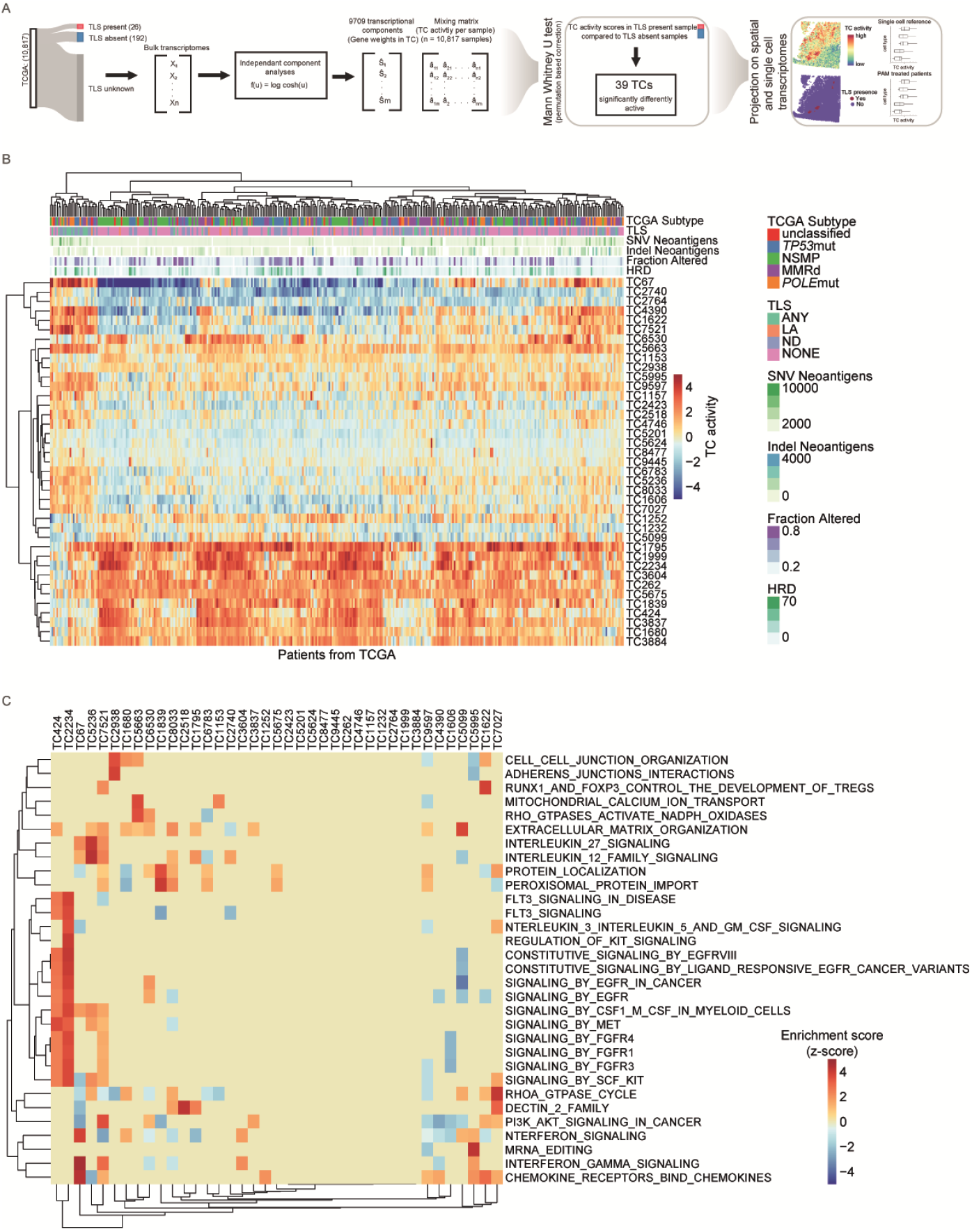
Transcriptional components associated with the presence of TLS. (A) Workflow for the data acquisition and consensus-independent component analysis (c-ICA). (B) Clinico-pathological variables (subtypes of cancer, presence of TLS, SNV neoantigens, Indel neoantigens, Fraction altered, Homologous recombination defects) per sample (top panel). Activity scores of the 39 TCs found to be associated with presence of TLS in Mann-Whitney U test (bottom panel). Clustering of samples was performed using Euclidean distance and the complete method, with heatmap colors based on activity scores. (C) Enrichment heatmap of Reactome gene sets in each TC associated with presence of TLS. GSEA results of 39 TCs associated with presence of TLS in Mann-whitney U test are presented, including Reactome gene sets that were included in the enrichment for at least one TC that passed the Bonferroni threshold for multiple testing correction. The gene sets were clustered using Euclidean distance and the complete method, and the heatmap colors were based on Z-scores, truncated at a value of five.

Gene-set enrichment analysis, using 12 gene-set collections from the molecular Signature DataBase (mSigDB)^42,43^, revealed clear enrichment of distinct biological processes within the TLS-associated TCs (Figure 5C, Source Data). The z-scores representing the enrichment of the most significant gene-sets per TLS-associated TCs ranged from 2.84 to 10.91 (IQR 3.93-5.1; median 4.35). Cross-study projection of these TLS-associated TCs onto the spatial transcriptomic profiles revealed that TC4390 was highly active in the EC TLS regions (Figure 6A-C). In line with our earlier results, TC4390 showed strong activity in Tex cells (Figure 6D and S15). Notably, the TC with the highest association with TLS presence in bulk gene expression profiles, TC1622, was also associated with a subset of Treg cells, showing high enrichment of gene-set; RUNX1 and FOXP3 controlling the development of Treg cells. The genes with the highest weights in TC1622 were indeed *FOXP3, FCRL3* and other genes related to T reg biology such as *CTLA-4, CCR4, CCR8, ICOS* and *TIGIT*. Finally, we explored the potential co-functionality of the genes with the highest weights in TC4390 using guilt-by-association analysis with gene set for mammalian phenotypes^44^(Figure 6E). We identified two clear co-functionality networks: one associated with abnormal B cell function and reduced CD8 T cells, and another related to embryonal/prenatal lethality, the latter likely reflecting the essential role of some TLS-related processes in development. Notably, the network related to B cell physiology not only captured genes known to be involved in TLS-GC regulation, such as *CXCL13, CR2* (CD21), and *FCRLA*, but also included genes not previously linked to TLS, such as *P2RY10* (encoding P2Y10) and *NAMPT. P2Y10* was recently reported to promote chemokine receptor-mediated migration of CD4 T cells^45^, while *NAMPT* was originally identified as a cytokine that synergizes with IL-7 and stem cell factor (SCF) in colony formation of pre-B cells^46^.

**Figure 6.**
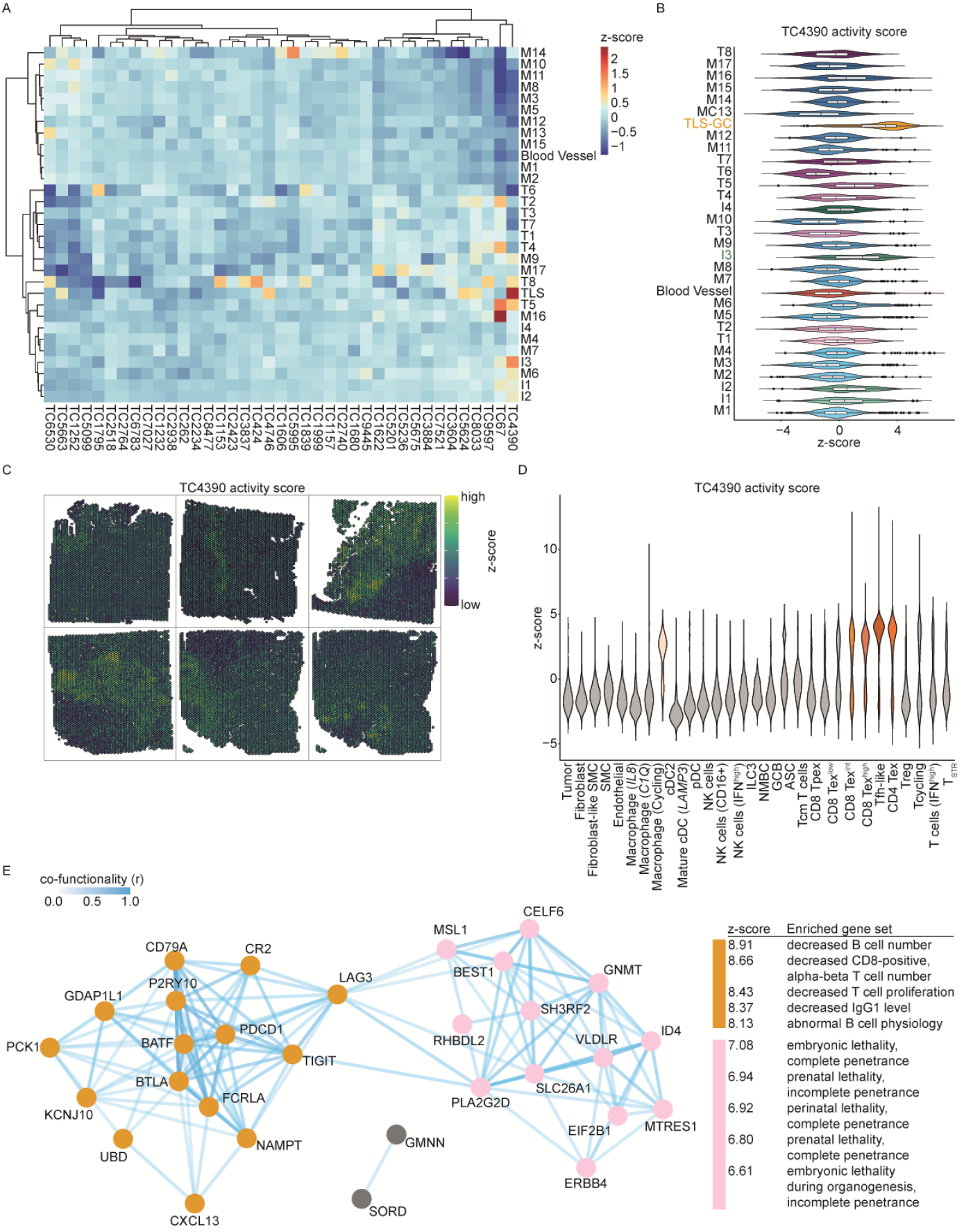
Characterization of TLS-associated TC4390. (A) Heatmap displaying the activity scores of 39 TCs across different ST clusters. (B) Violin plot depicting the activity score of TC4390 across various ST clusters. (C) Spatial expression pattern of TC4390 in different EC samples. (D) Violin plot illustrating the activity score of TC4390 across all immune cell types. (E) Co-functionality network of genes associated with TC4390, along with the biological processes of these genes.

These, and other genes from TC4390, therefore provide an excellent starting point for further functional dissection of TLSs.

### Transcriptional patterns of TLS during neoadjuvant ICI in EC patients

Having established the conserved TC that spatially localizes to TLSs in EC, we next sought to understand how the activity of this TC is altered during anti-PD1 immunotherapy, which is clinically associated with the induction of TLS. To this end, we analyzed a recently published cohort of T cell scRNAseq data from three MMRd EC patients treated with two cycles of neoadjuvant ICI^47^. Data on ICI-induced T cell clonal expansion was available for all T cells in this patient cohort, as was information on clonal sharing with the TDLN. We used the original phenotype classification from the study and further categorized cells into four groups: tumor-infiltrating T cells from ICI-responsive and ICI-nonresponsive patients as well as T cells from TDLN of ICI-responsive or ICI-nonresponsive patients. Cross-study projection of TLS-TCs revealed higher activity of TC4390 in exhausted CD8 T cells, exhausted and effector CD4 T cells, and cycling T cells (Figure 7A and S16). The only other TC significantly active within this dataset, TC1622, was active in Treg cells (Figure S16), which is consistent with our findings in scRNAseq projection (Figure S15). Further analysis revealed specific and significant activity of TC4390 in ICI-responsive T cells, both in TDLN- and non-TDLN-associated single cells (Figure 7B). The activity of TLS-associated TC4390 was not directly correlated with the overall clone size or fold expansion. Both smaller and larger clones exhibited a near-uniform high, low, or intermediate TC4390 activity (Figure 7C). This pattern held true for both TDLN (Figure 7C) and non-TDLN-associated clones (Figure 7C).

**Figure 7.**
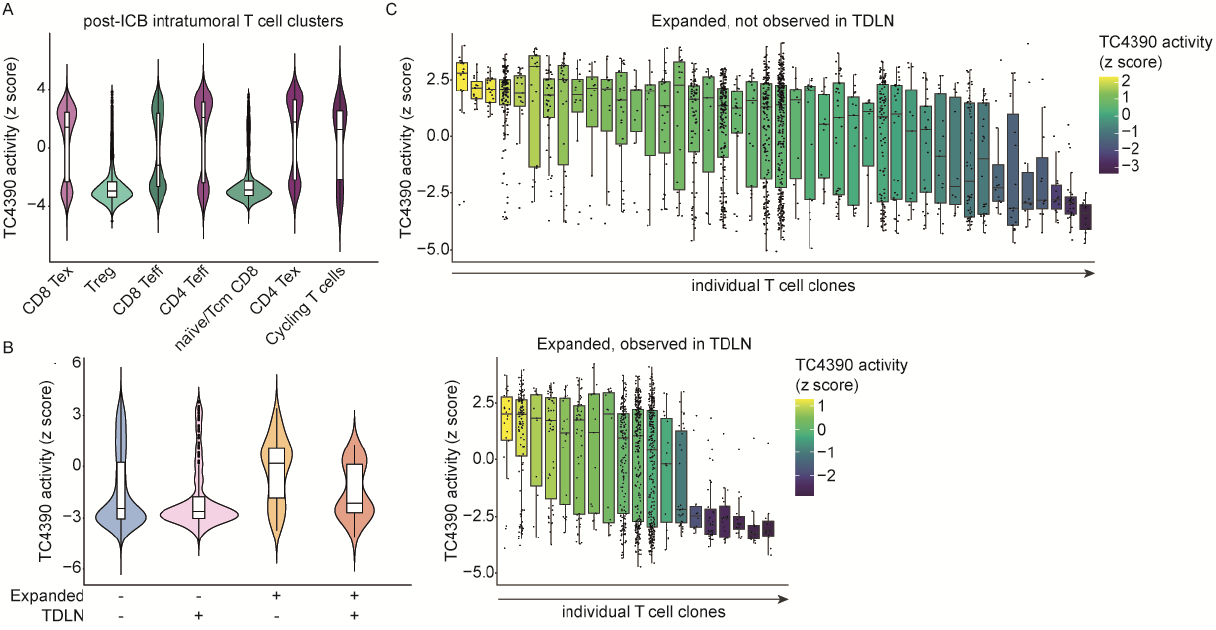
TC4390 activity upon ICI treatment. (A) Violin plot depicting z-score of TC4390 activity across intra-tumoral T cell clusters in EC-patients post ICI-treatment. (B) Violon plot depicting z-core of TC4390 across both TDLN and non-TDLN-associated single cells. (C) Box-and-whisker plots of TC4390 activity across individual T-cell clones both in TDLN- and non-TDLN-associated T-cell clones.

In summary, we conclude that neoadjuvant ICI induces the activity of the transcriptional component significantly linked to TLS in bulk and spatial gene expression profiles, potentially providing a mechanistic basis for TLS induction.

## DISCUSSION

In the present study, we leverage harmonized multi-level transcriptomics to comprehensively define the spatial architecture and immunobiology of TLSs and their contribution to anti-tumor immunity in EC. We reveal the organizing principles of TLSs through chemokine gradients, the dissemination of class-switched B cells from TLS, and a conserved transcriptomic pattern for TLS across hundreds of EC tumors. Additionally, we demonstrate how ICI induces the activity of this transcriptomic pattern associated with TLS formation.

Our study sheds new light on several aspects of TLS neogenesis. It has been established that post-capillary vasculature in chronically-inflamed tissue, such as immunogenic cancers, can recruit activated lymphocytes and APCs^48–50^. However, the subsequent NF-κB-mediated differentiation of postcapillary venules into HEVs, which express peripheral node addressin (PNAd), is required for the recruitment of naive (CD62L+) lymphocytes and the formation of mTLS^51,52^. In this study, we show that these HEV-containing TLSs are spatially localized near peripheral *CCL21*+ LECs, which closely resemble the cortical and paracortical LECs juxtaposed to HEV in lymph nodes. We propose that the dense capillary/lymphatic network of the myometrium, where EC-associated TLSs are almost invariably localized^17^, serves as a major site for the ingress and egress of immune cells from the tumor bed. It is reasonable to suggest that the progressive spatial accumulation of TNFα, LTα, and/or LTαβ expressing immune cells in the myometrium drives NF-κB signaling in endothelial cells, which is required for HEV formation^51^. Supporting this, (re)circulating LTαβ-expressing DCs are known to maintain the HEV network in lymph nodes of the adult mouse^53,54^, a mechanism that may similarly apply to APCs during the immune surveillance of a developing tumor. Since CCL21 induces LTα1β2 expression on naive CD4 T cells^55^, the formation of HEC cells and recruitment of naive T cells likely represents a self-amplifying process in the capillary bed. Several lines of evidence also point to perivascular cells as precursors of ectopic follicular reticular cells (FRCs) and FDCs^56–58^. As FRC/FDC differentiation is driven by LTβR signaling^57^, and FRCs are present in the perivascular sheath surrounding HEVs^51^, it is plausible that differentiation of these key TLS constituents occurs in tandem.

Our data, along with others^33^, suggest that at a certain point in tumor development, mDC “fail” to migrate into the TDLN and remain within the tumor. Increased local production of CCL21 by FDCs may be a key determinant in “routing” mDC away from draining peripheral LECs and toward TLSs, where DC-produced cytokines help recruit and organize Tfh, Treg cells, and CXCL13+ Tex cells. Our spatial colocalization data supports an interaction between mDC and *CXCL13*+ CD4 T cells in the T cell zone of TLS, as evidenced by the high expression of a DC help signature in mDC. In the context of immunotherapy, intratumoral niches of physically interacting CXCL13+ CD4 T cells and mature LAMP3+ DCs have been described in lung and liver cancer^59,60^. These CD4/LAMP3+ DCs hubs included progenitor CD8+ T cells, suggesting that these triads control the differentiation of tumor-specific CD8. TLSs in our study exhibited similar immune cell interaction patterns, though we were unable to pinpoint Tpex cells in our spatial analysis, potentially due to their rarity. Our data further supports the notion that TLSs serve as sites for the development of adaptive anti-tumor immunity *in situ*. Although we could not perform a proper clonal analysis of the TCR due to the nature of the FFPE tissue, we did observe increased TCR diversity at the V gene segment level when compared to more distal regions.

Consistent with previous reports, we also found evidence of local B cell differentiation in TLSs, from NMB cells to class-switched ASCs, which disseminate into the TME^29^. The levels of scattered IgA/IgG, along with the inverse correlation between IgD/IgM and IgG/IgA from the TLS center, indicates affinity maturation of B cells and potential antigen-specific interactions within TLSs. Although we could not determine the tumor specificity of the antibodies produced by the B cells in this study, the spatial colocalization of *IGHA1*+ ASCs with its cognate receptor PIGR suggests that non-antigen-specific interactions are likely to occur. PIGR promotes activation of tumor-cell intrinsic^61^ and extrinsic immune responses following binding of transcytosed secreted IgA (sIgA)^26,27,62^, and intratumoral IgA levels have been correlated with improved survival in EC^61^. Additionally, sIgA has been shown to engage FCRL3 expressed on FOXP3+ Treg cells, inhibiting their suppressive function and promoting a transition towards an IFN-γ-expressing Th17 phenotype^28^. In our study, FCRL3+ cells were distinctively localized around TLSs, and ICA analysis revealed a strong correlation between TLSs and activity of a TC comprising *FCRL3* and *FOXP3* as top genes.

Given the presence of *FCRL3*+ Treg cells and the abundance of IgA surrounding TLS, sIgA-FCRL3 interactions may inhibit immunosuppressive Treg cell function and promote an IFN-γ-producing anti-tumor phenotype^28^. Anti-tumor Treg cells have been previously described in colorectal cancer, head and neck squamous cell carcinoma, and gastric cancer^63^. Conversely, Treg cells, particularly FCRL3-expressing T regs, are known to support tumor growth and have been shown to be highly immunosuppressive^64^. In EC, both the presence of intra-tumoral Treg cells and the CD8+/T reg ratio have been associated with higher tumor grade and worse prognosis^65–67^. In mouse models of lung adenocarcinoma, Treg cells have been shown to actively suppress anti-tumor T cell immunity inside TLSs^68^, and the presence of Treg cells in or around TLS correlates with worse prognosis in patients with lung cancer^69^. While the immunosuppressive functions of Treg cells are well recognized, their presence within TLS in our study does not necessarily imply an immunosuppressive role. It is plausible that their accumulation within these structures is a default biological response to counteract excessive inflammation, as TLS are indicative of heightened immunogenicity^7,17^. Thus, whether FCRL3+ Treg cells around TLS are merely bystanders, or whether they exert anti- or pro-tumor functions in EC, remains to be addressed.

Most transcriptomics studies, including those analyzing TLS, used either spatial or single cell RNA-seq with small sample sizes or relied on bulk transcriptomes. While bulk transcriptome datasets are usually large, the profiles are generated from samples that contain tumor, stromal and immune cell components, reflecting the average transcriptional patterns of all biological processes present in the tumor samples. Subtle transcriptional patterns related to processes, such as TLS formation, are often “overshadowed” by dominant patterns from other biological processes. C-ICA offers an approach to harmonize all three platforms by decomposing bulk transcriptomes into statistically independent TCs and projecting them onto spatial and scRNAseq data. As demonstrated here, this approach allowed us to identify the unique TCs associated with TLS in bulk transcriptomes, spatially co-localized with TLS, and significantly active in single cell transcriptomes. This approach also enabled us to project these TCs onto an independent dataset from patients treated with neoadjuvant ICI. The results showed how ICI-responsive clones had increased activity of the TC spatially-associated with TLS, potentially providing a mechanistic basis for how ICI might induce TLS. Gene-level analysis of the TLS-TCs also revealed novel genes, such as chemokines, previously implicated in immune biology, but not yet directly linked to TLS, which warrant further investigation. Future work will need to address how these TLS-TCs spatially alter upon ICI as more longitudinal on-treatment ICI-datasets become available.

Finally, to the best of our knowledge, this is the first study to apply spatial transcriptomics in EC. While we specifically focused on the immune composition and interactions in the TME, this dataset can be further leveraged to study the complex, multi-dimensional cellular and molecular landscape of EC cells and their interaction with the TME.

Altogether, our study provides novel insight into the organization of TLSs, offers a potential mechanistic basis for ICI-induced TLS formation and identifies novel gene- and TME-level targets for further mechanistic and therapeutic exploration.

## ACKNOWLEDGMENTS

We thank all the patients, and their families for their contributions. The present study was funded by a UMCG Cancer Research Fund grant to M.R., M.B, H.W.N. and R.F.

## AUTHOR CONTRIBUTIONS

M.R., K.B., A.B. collected, analyzed, and interpreted the data and wrote the manuscript with input, edits and approval from all authors. M.R., N.R. and A.P. performed spatial transcriptomics and single-cell RNA-sequencing experiments. A.B. performed the ICA analysis. F.S. and D.H. contributed to the analysis of the data. A.V. collected patient samples. A.B., R.W., D.S., F.F., and M.B. performed bioinformatics analysis. T.B. and J.B. assessed tumor tissue and performed IHC staining and molecular profiling. M.B., R.F. and H.W.N. conceived the study, wrote the study protocol, analyzed, interpreted the data, and supervised writing of the manuscript.

## DECLARATION OF AI AND AI-ASSISTED TECHNOLOGIES

This manuscript utilized artificial intelligence (AI) and AI-assisted technologies to assist in rewriting specific passages of text for clarity and conciseness. The AI tool, ChatGPT (OpenAI), was employed to rephrase sentences and improve readability without altering the scientific content or meaning. The authors carefully reviewed all AI-generated suggestions to ensure accuracy, appropriateness, and alignment with the manuscript’s scientific objectives. No AI algorithms were used for data analysis, generation of original research content, or drawing scientific conclusions. The AI assistance was solely employed as a writing aid to enhance the clarity of the text. The use of AI tools in manuscript preparation complies with ethical guidelines and policies set forth by the journal.

## DATA AVAILIBILITY STATEMENT

The raw and processed scRNAseq data discussed in this publication have been deposited in NCBI’s Gene Expression Omnibus and are accessible through GEO Series accession number GSE^******^. The clinical data, and tumor material generated during, or analysed in, the present study are not publicly available owing restrictions by privacy laws. Data are held by the coauthors of this article. Requests for sharing of data and material should be addressed to the corresponding author(s) within 15 years of the date of publication of this article and include a scientific proposal. Depending on the specific research proposal, the coauthors will determine when, for how long, for which specific purposes and under which conditions the requested data can be made available, subject to ethical consent and the composition of a material and/or data transfer agreement. Requests for data access will be processed within a 3-month timeframe. The remaining data are available within the publication, Supplementary information or Source Data.

## CODE AVAILIBILITY STATEMENT

The complete set of codes utilized in this study is available at a github repository: https://github.com/arkajyotibhattacharya/TLSLandscapeinEndometrialCancer

## SUPPLEMENTARY FIGURE LEGENDS

**Figure S1.**
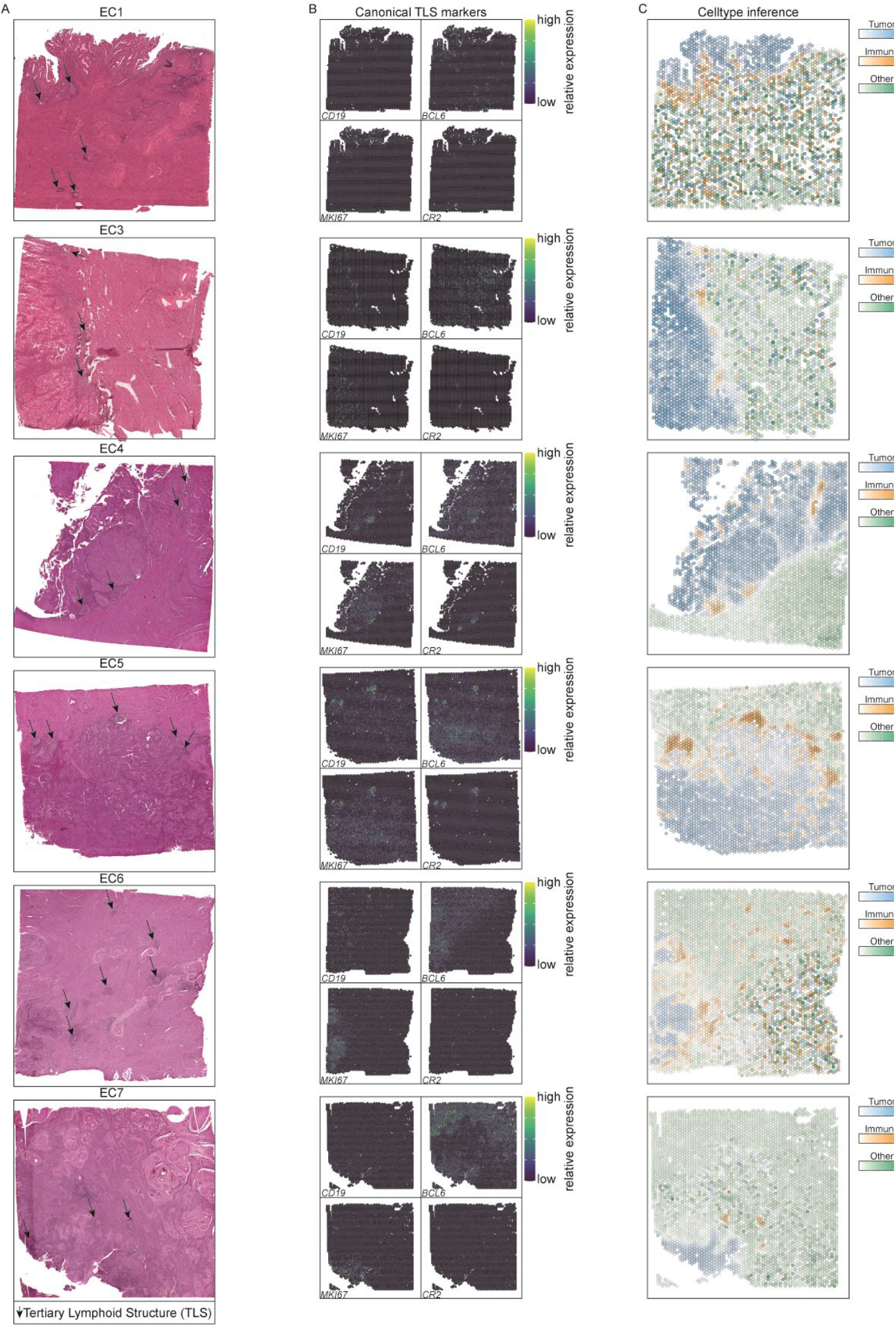
Overview of the EC samples used. (A) H&E staining of all EC sections in which ST was performed. Black arrows indicate the pathologist’s annotation of TLSs. (B) Spatial gene expression of *CD19, BCL6, MKI67* and *CR2* across all EC patient sections used in this study. (C) Celltype inference depicting tumor (blue), immune (orange) and other (green) cell type ST spots across all EC patient sections used in this study.

**Figure S2.**
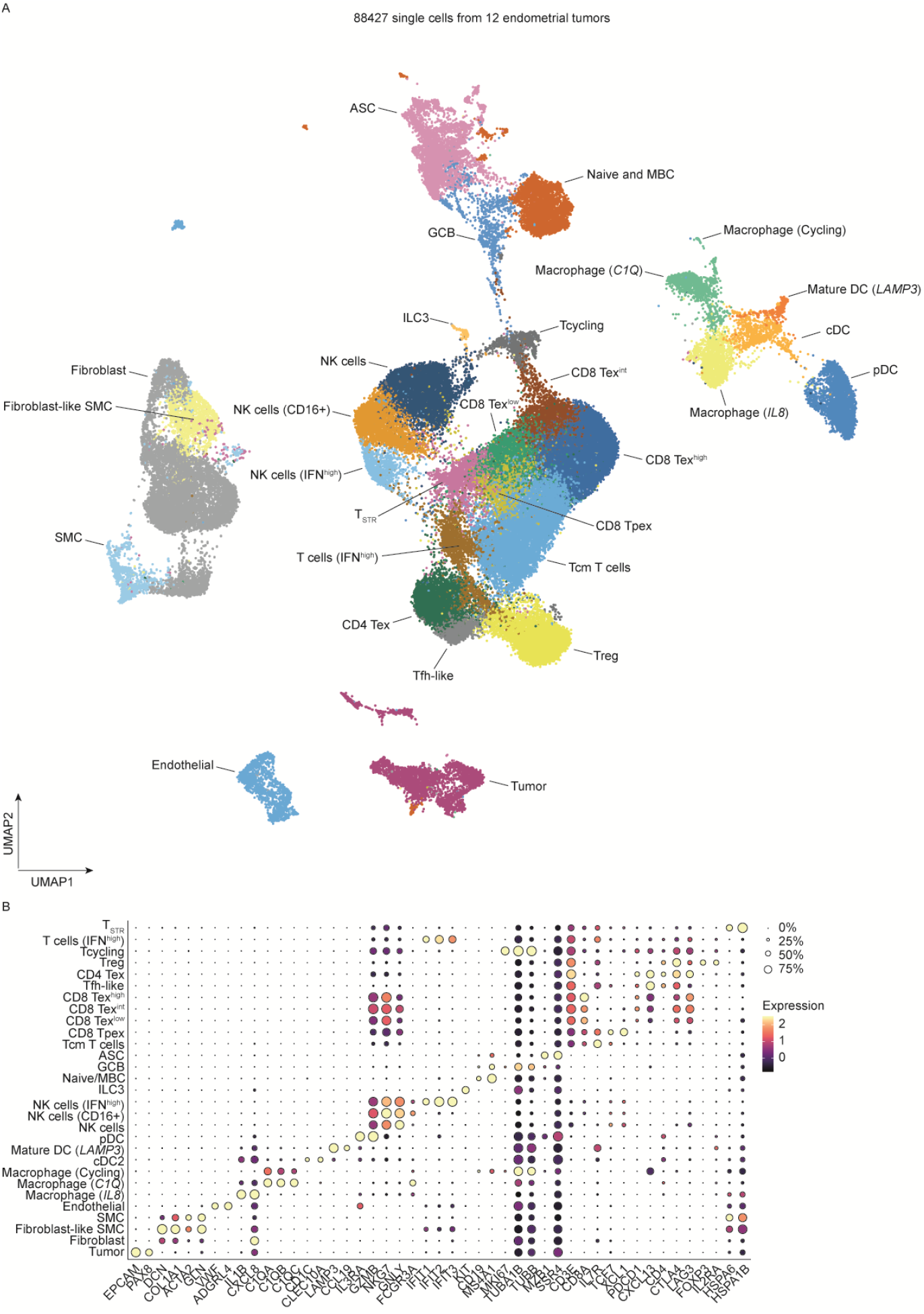
Single-cell atlas of EC. (A) UMAP visualization of integrated scRNAseq data from endometrial tumors, combining datasets from Regner et al. and our own in-house generated dataset. Clusters were identified by unsupervised hierarchical clustering. (B) Dotplot of canonical marker expression used for annotation of the major cell populations identified in A. Cell types were identified based on canonical marker expression and differential gene expression. CD8 Tex cells were classified in high, intermediate or low based on the gradient expression of *CXCL13, LAG3* and other differentially expressed genes. T_STR_: stressed T cells; T reg: regulatory T cells; T_ex_: exhausted T cells, T_fh_-like: follicular helper T cells, T_pex_: precursor-exhausted T cells; T_cm_: central-memory T cells; ASC: antibody-secreting cells; GCB: germinal-center B cells; Naïve/MBC: naïve and memory B cells; ILC3: innate-lymphoid cells group 3; NK: natural killer cells; pDC: plasmacytoid dendritic cells; SMC: smooth muscle cells.

**Figure S3.**
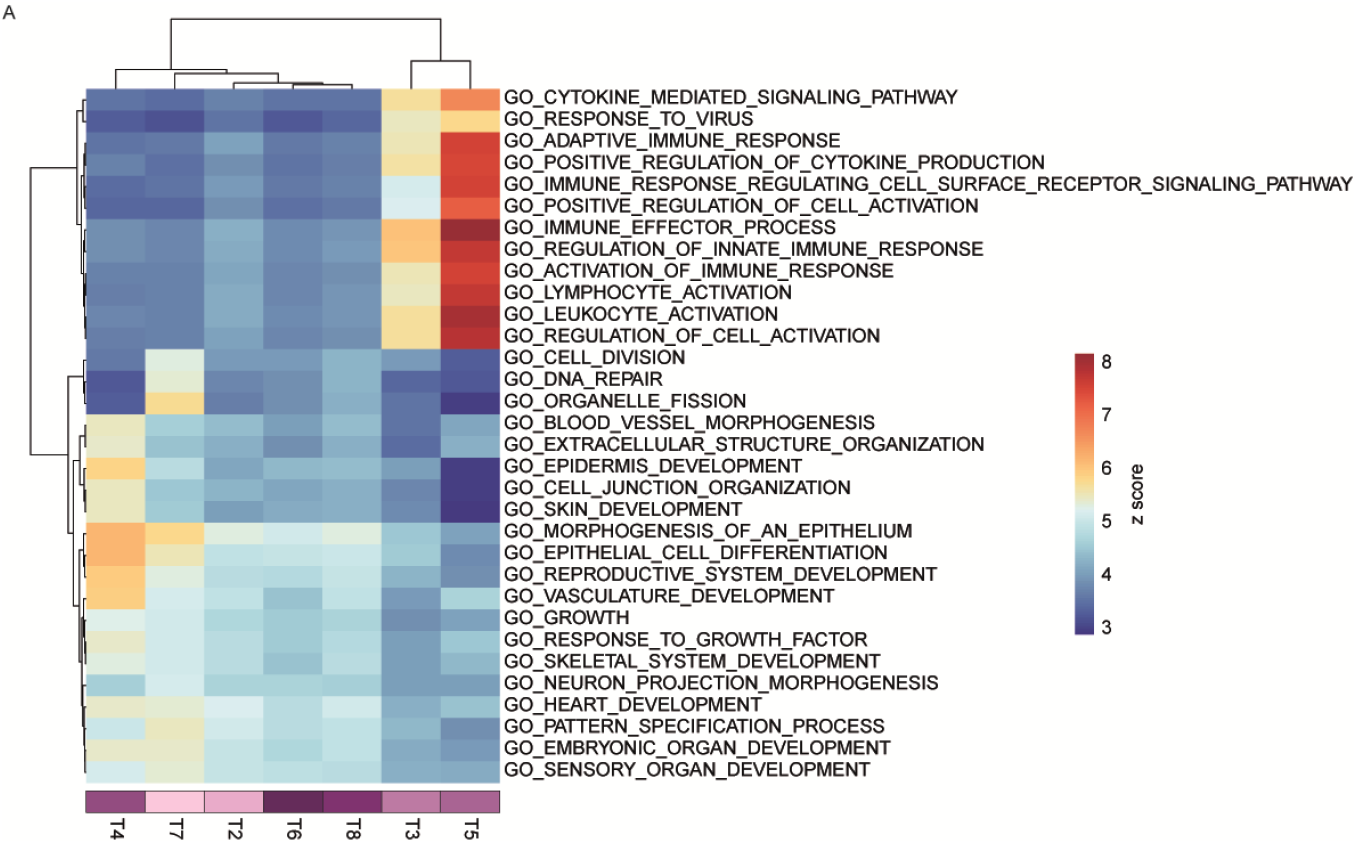
Differential pathway analysis across the different tumor clusters. (A) Gene-ontology signaling pathways enriched in the endometrial tumor clusters identified through guilt-by-association analysis. Tumor cluster 1 did not show any differentially expressed pathways. Clustering of samples was performed using the Ward.D2 method, with heatmap colors based on Z scores.

**Figure S4.**
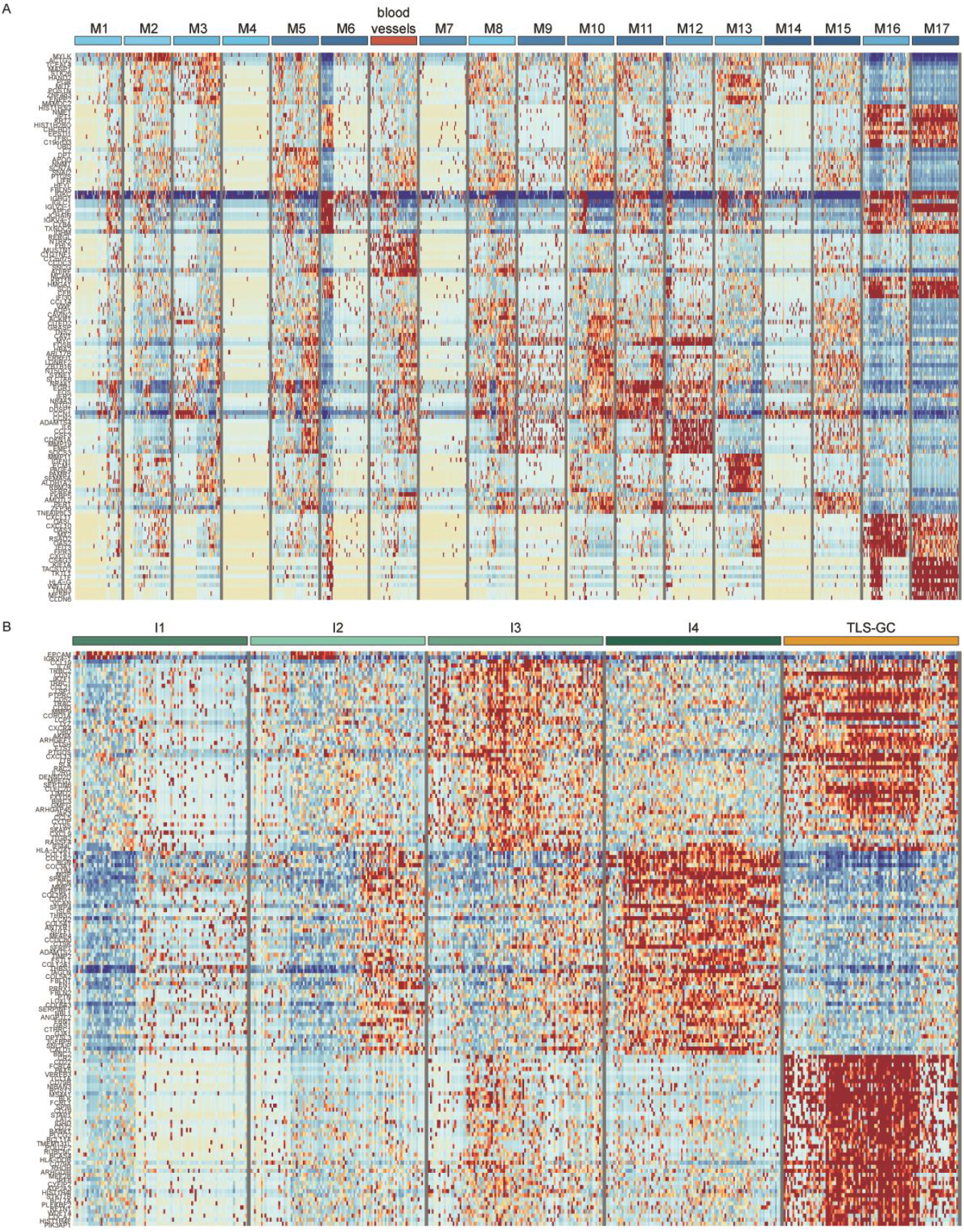
Characterization of myometrium and immune cell clusters in EC. Heatmap of the normalized and scaled differential gene expression values across (A) myometrium and blood vessel clusters and (B) immune cell clusters identified using spatial transcriptomics.

**Figure S5.**
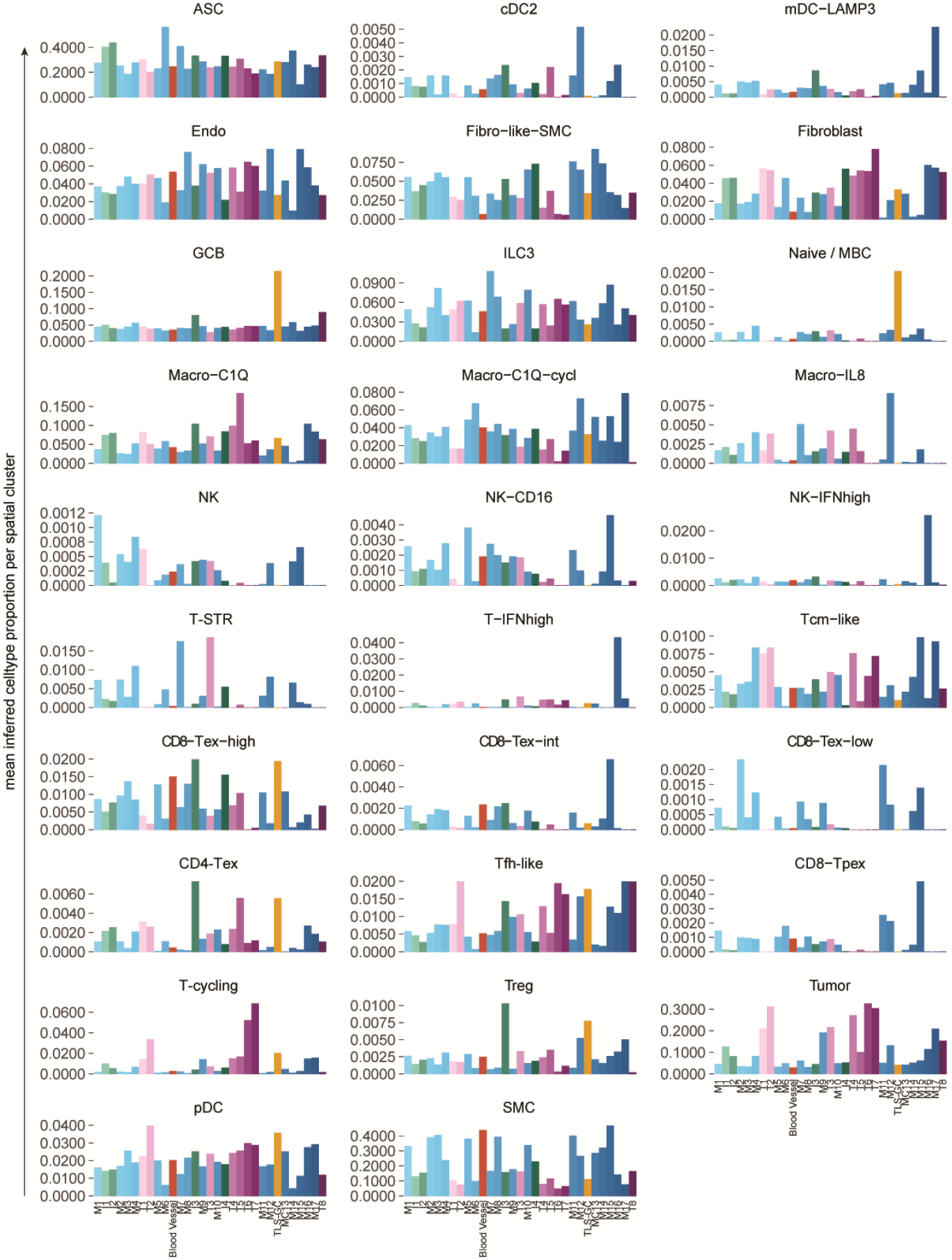
Cell type composition per spatial cluster. Mean relative inferred abundance of cell types for each defined spatial cluster annotated by tumor (purple), blood vessel (red), myometrium (bue), TLS (orange) and immune (green).

**Figure S6.**
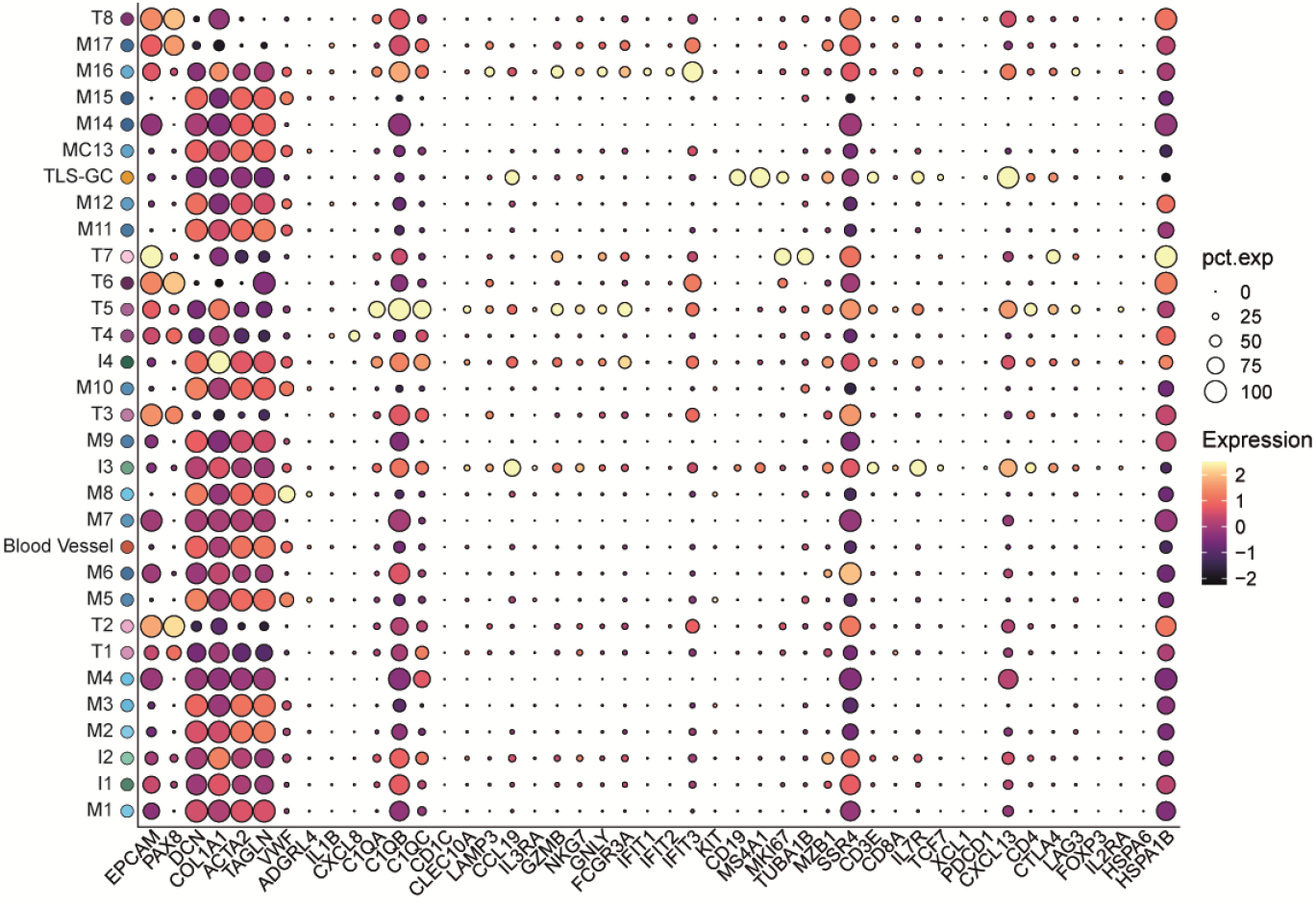
Canonical marker projection on spatial transcriptomics clusters. Dot plot displaying canonical marker expression used to annotate the single cell dataset, plotted for each major spatial cluster identified in the spatial transcriptomics dataset.

**Figure S7.**
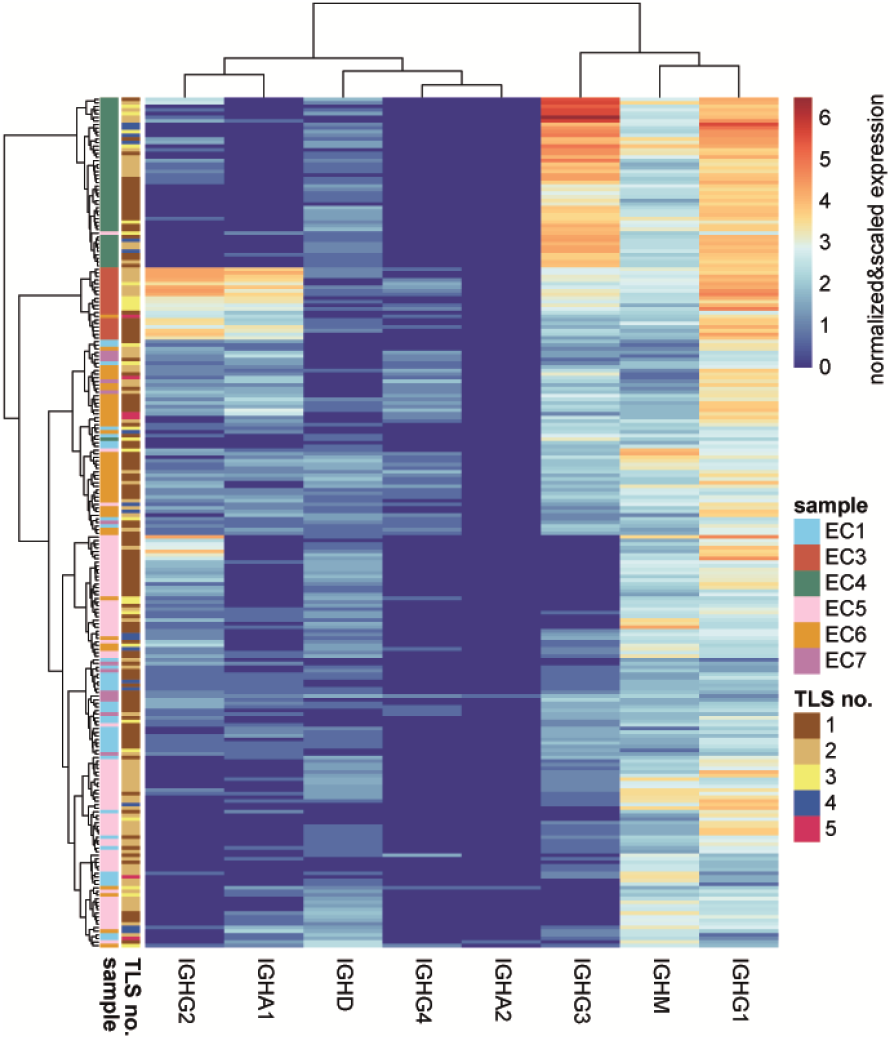
Distribution of immunoglobulin genes in TLSs. Heatmap illustrating the relative expression levels of *IgG, IgA, IgD* and *IgM* genes across individual TLSs.

**Figure S8.**
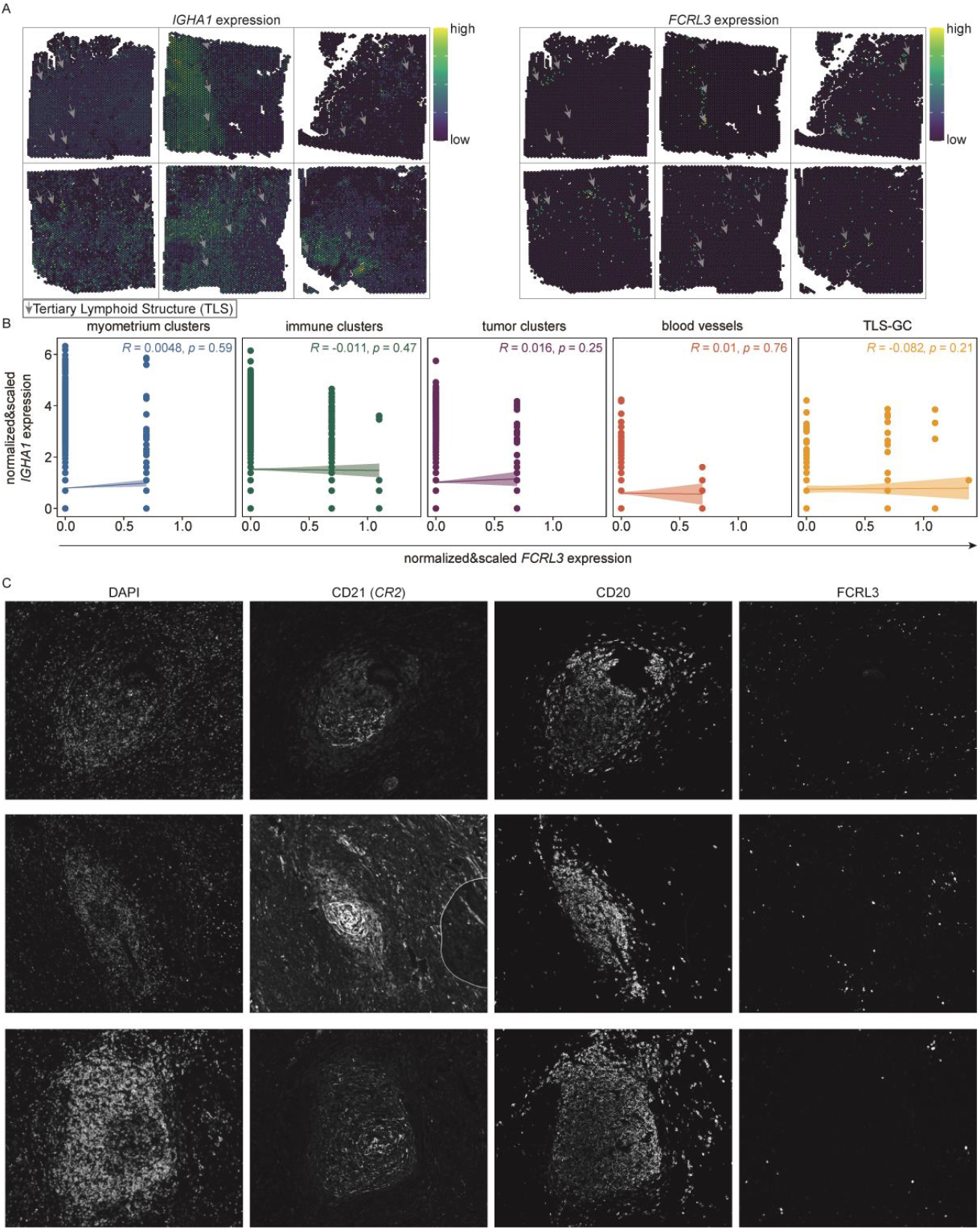
Expression of FCRL3 and IgA. (A) Spatial projections of *IGHA1* and *FCRL3* genes across the 6 EC samples. Grey arrows indicate the location of the TLSs. (B) Correlation plot between *IGHA1* and *FCRL3* across the main tissue regions identified by gene-level clustering and pathology annotation based on their normalized and scaled expression. (C) IF CD21, CD20 and FCRL3 staining of TLSs.

**Figure S9.**
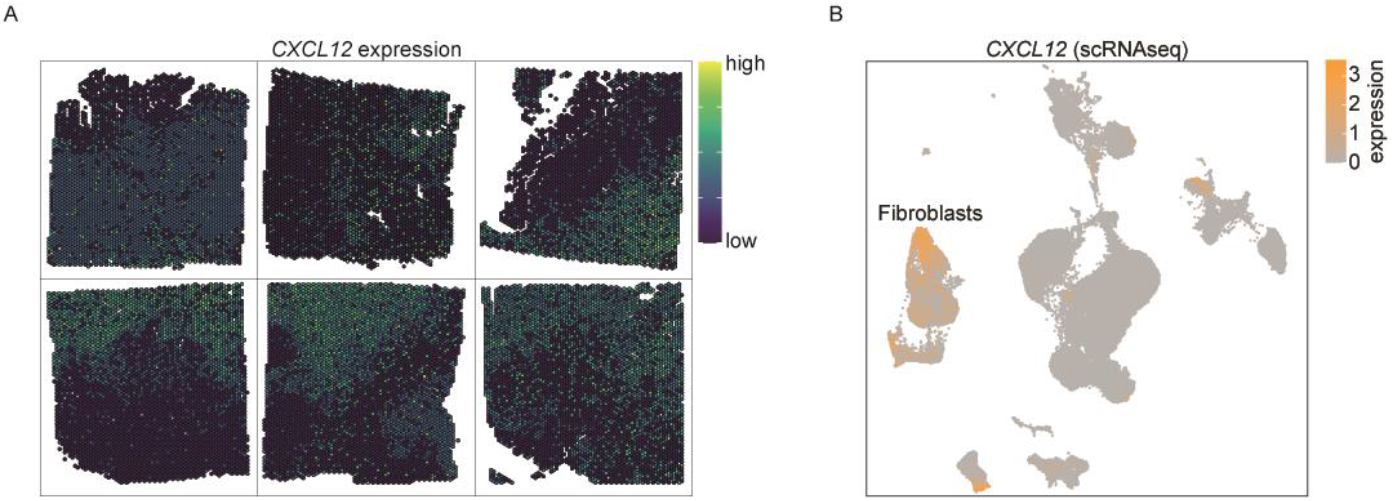
CXCL12 expression patterns. (A) Spatial projections of *CXCL12* expression across EC samples. (B) UMAP highlighting CXCL12 gene expression across clusters.

**Figure S10.**
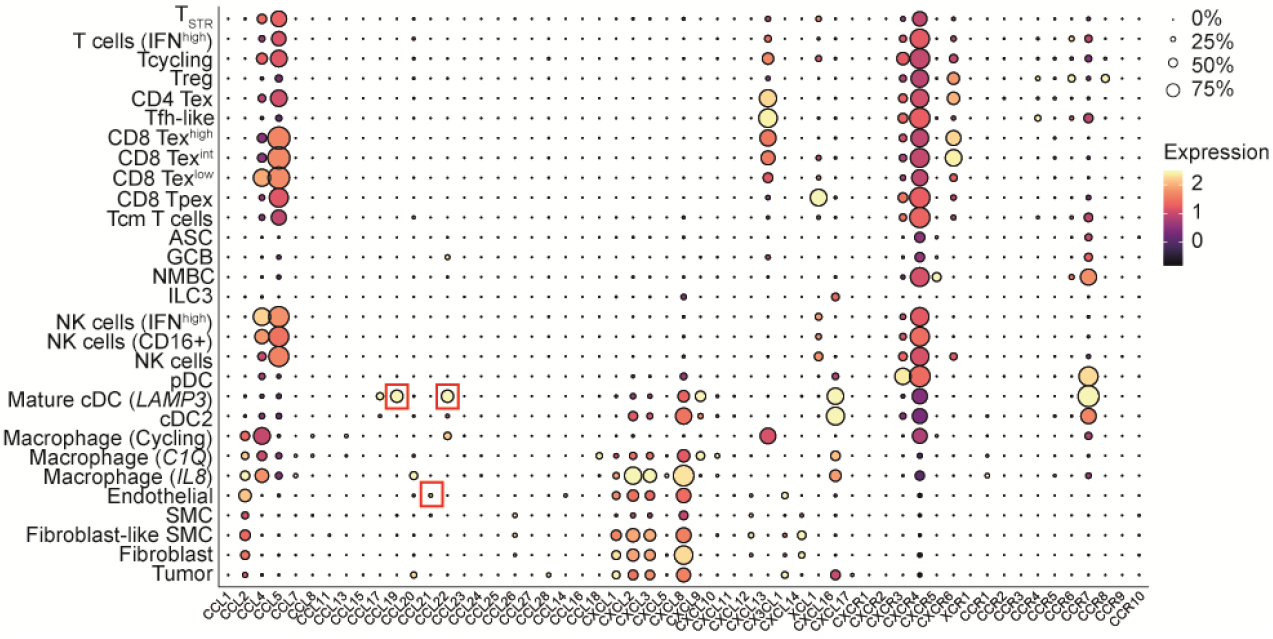
Chemokine landscape in TLS-positive EC samples. Chemokine and chemokine receptor gene expression levels for each cell subtype identified in the scRNAseq dataset.

**Figure S11.**
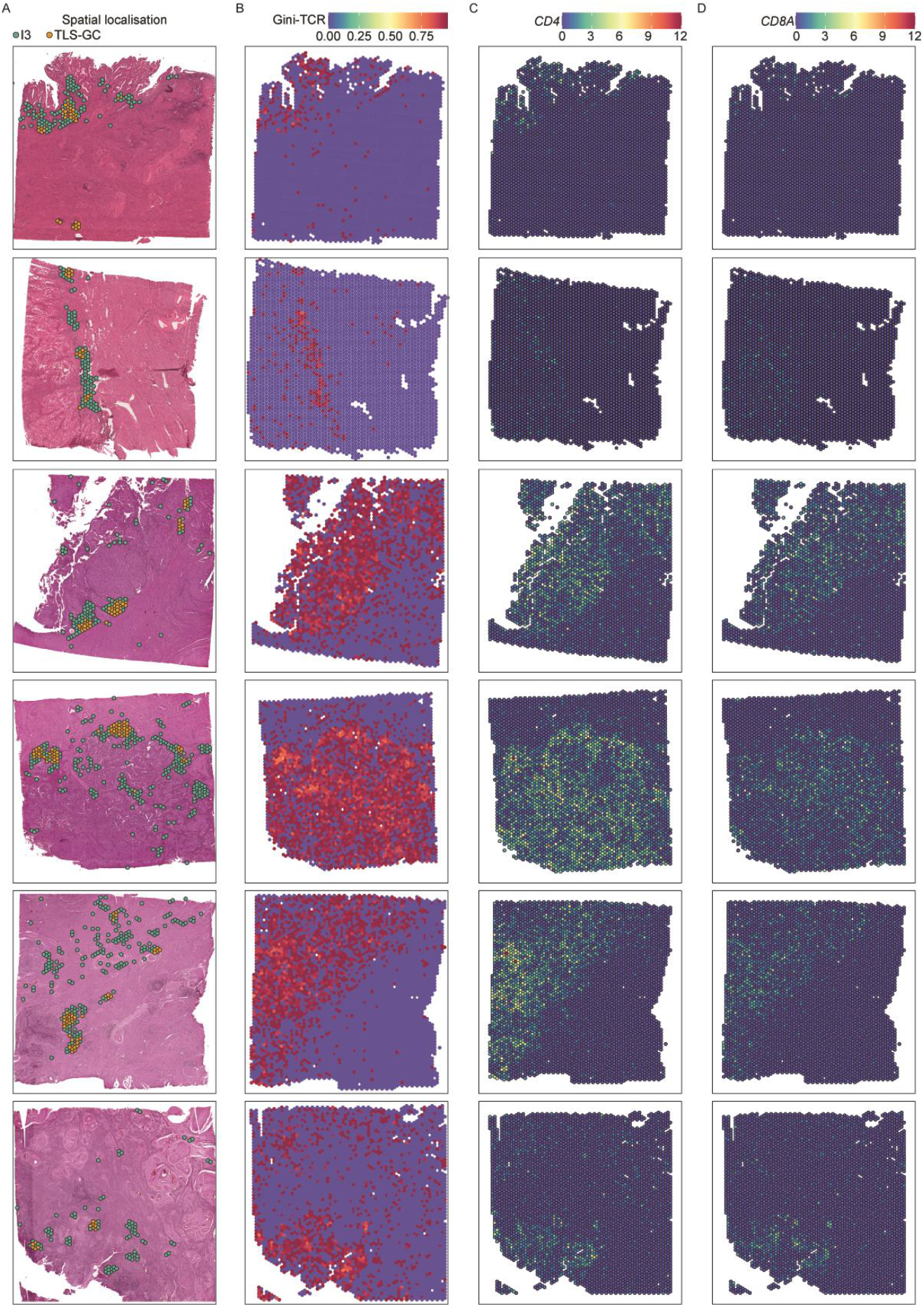
T cell composition of EC samples. (A) Spatial colocalization of immune cluster 3 (green) and TLS cluster (orange) across all EC samples projected on the corresponding H&E image. (B) Gini-TCR index across all EC samples. (C) Spatial projection of *CD8A* and *CD4* expression across all EC samples.

**Figure S12.**
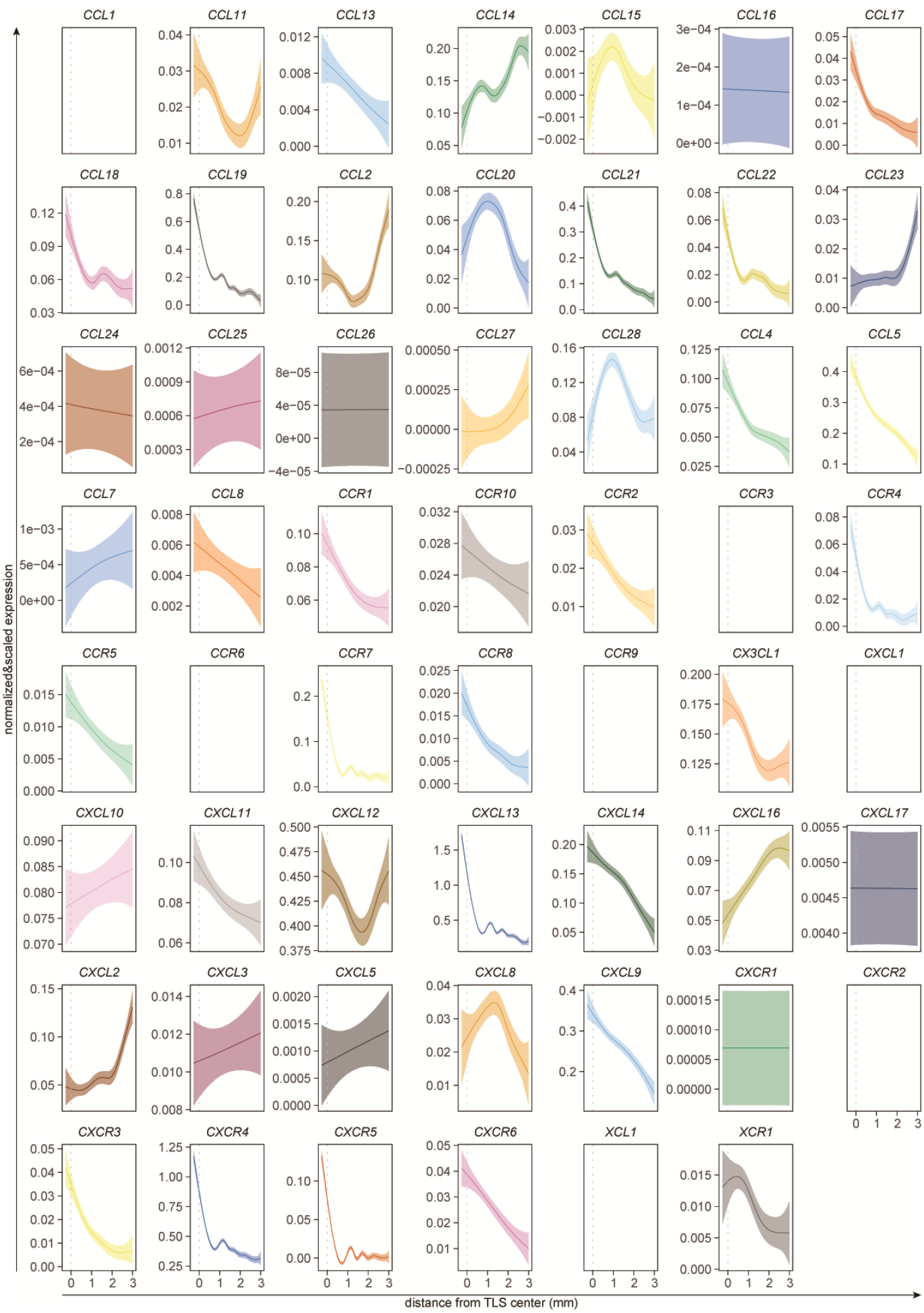
Chemokine gradients relative to TLS center. Chemokine and chemokine gene receptor expression expressed as radial distance using the TLS center as starting point.

**Figure S13.**
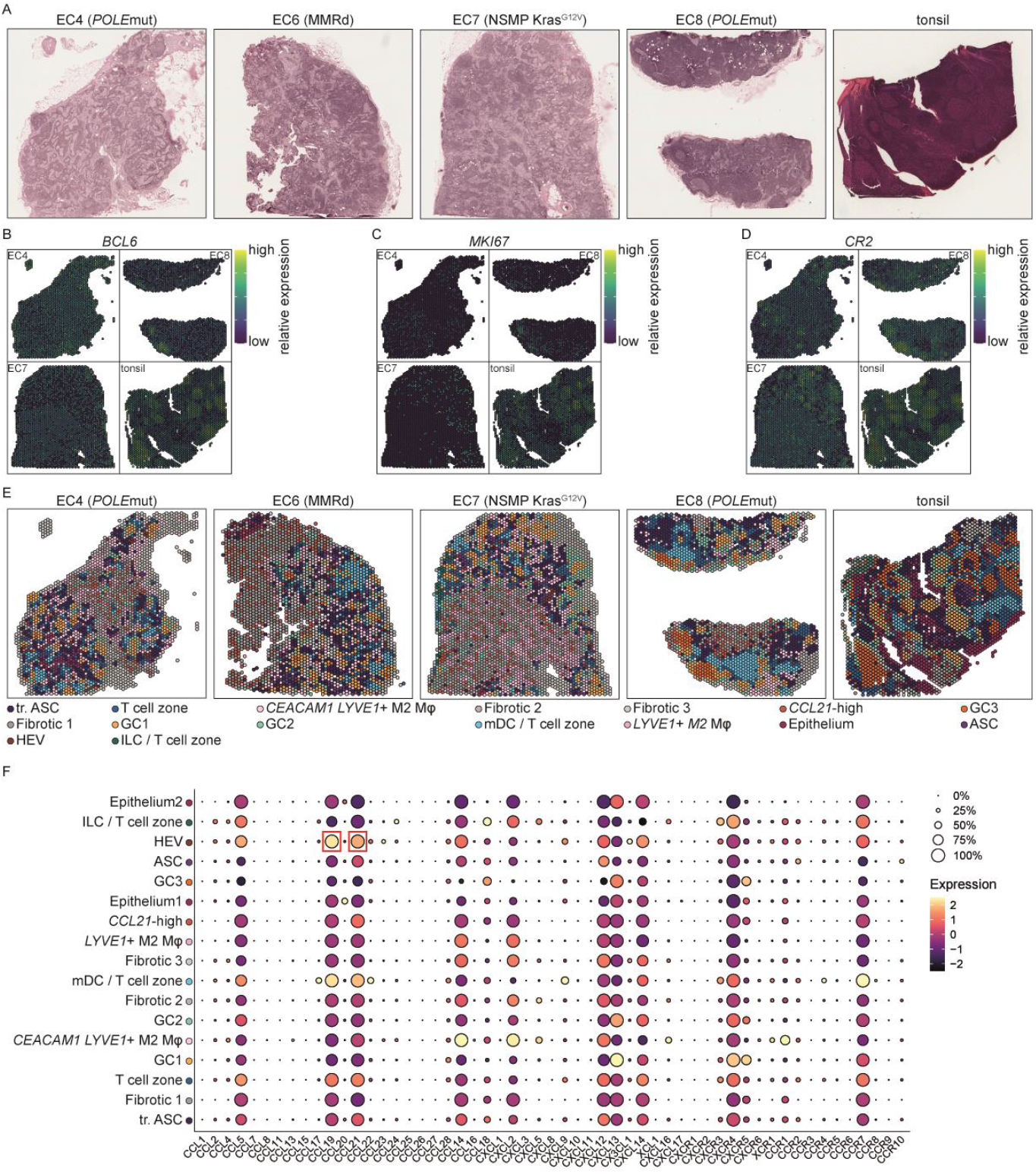
Landscape of TDLN and tonsil. (A) H&E images of all the TDLN and the tonsil reference used for spatial transcriptomics. Projection of the relative expression of (B) *BCL6*, (C) *MKI67* and (D) *CR2*. (E) Clusters of the ST spots across the TDLNs and the tonsil. (F) Chemokine and chemokine receptor expression levels for each ST cluster identified in the TDLN and tonsil. ASC: antibody-secreting cells; tr. ASC: transitioning antibody-secreting cells; HEV: high endothelial venule; GC1: germinal center 1; ILC: innate-lymphoid cells; GC2: germinal center 2; mDC: mature dendritic cells; GC3: germinal center 3.

**Figure S14.**
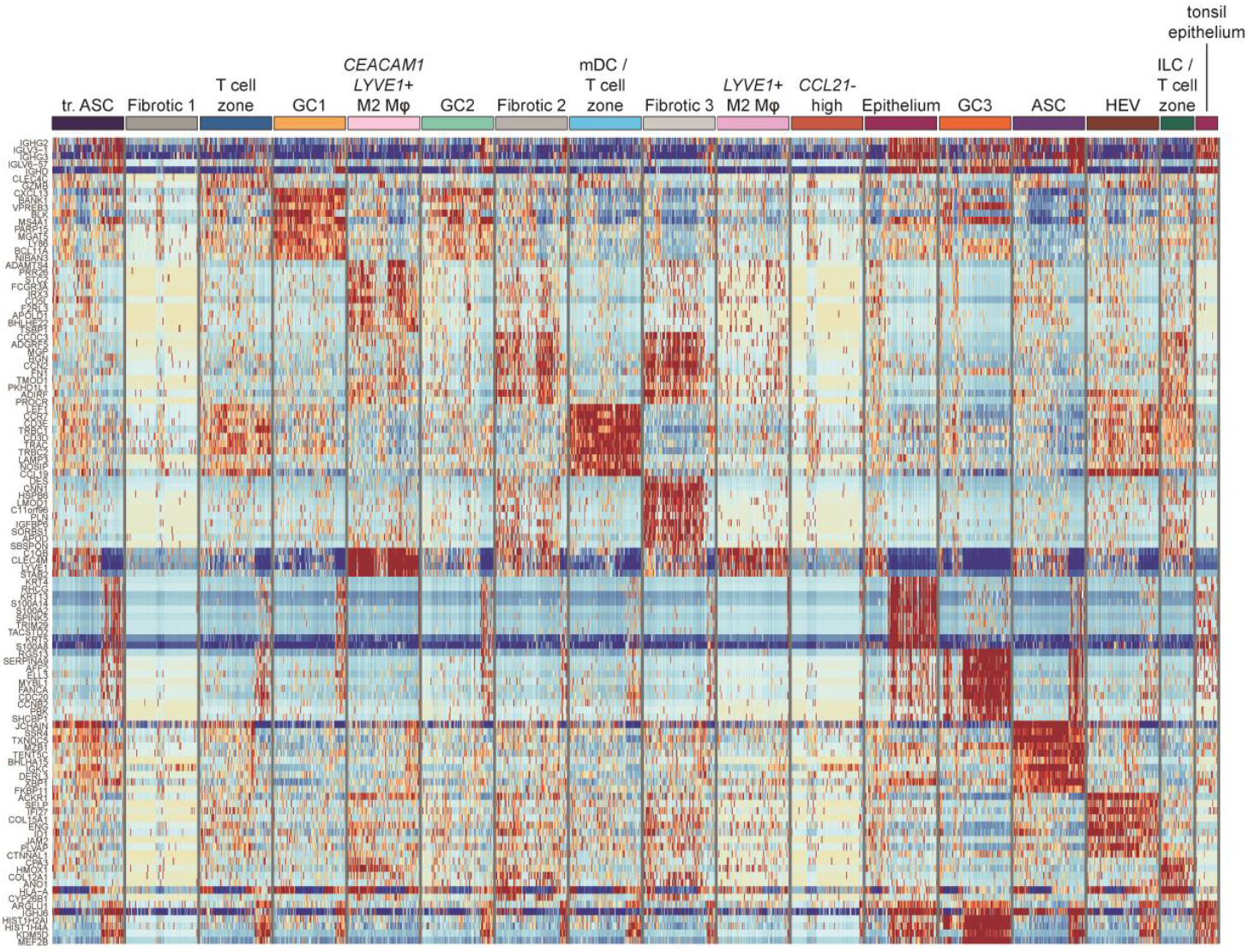
Characterization of immune cell clusters in the TDLNs. Heatmap of the normalized and scaled differential gene expression values across the TDLN clusters.

**Figure S15.**
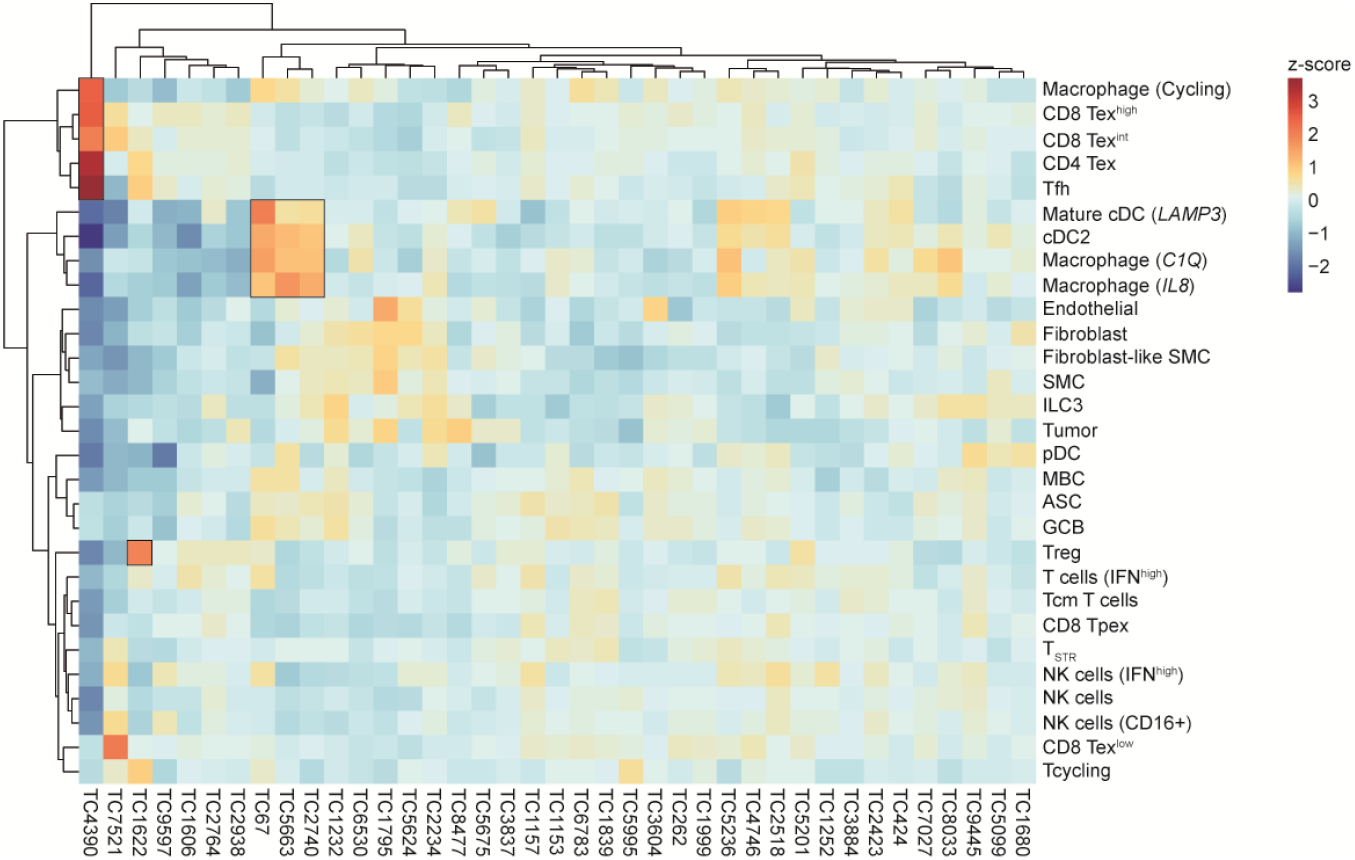
Activity of TLS-associated TCs across immune cell subsets. Heatmap displaying the activity scores of 39 TCs across the different clusters identified in the scRNAseq dataset.

**Figure S16.**
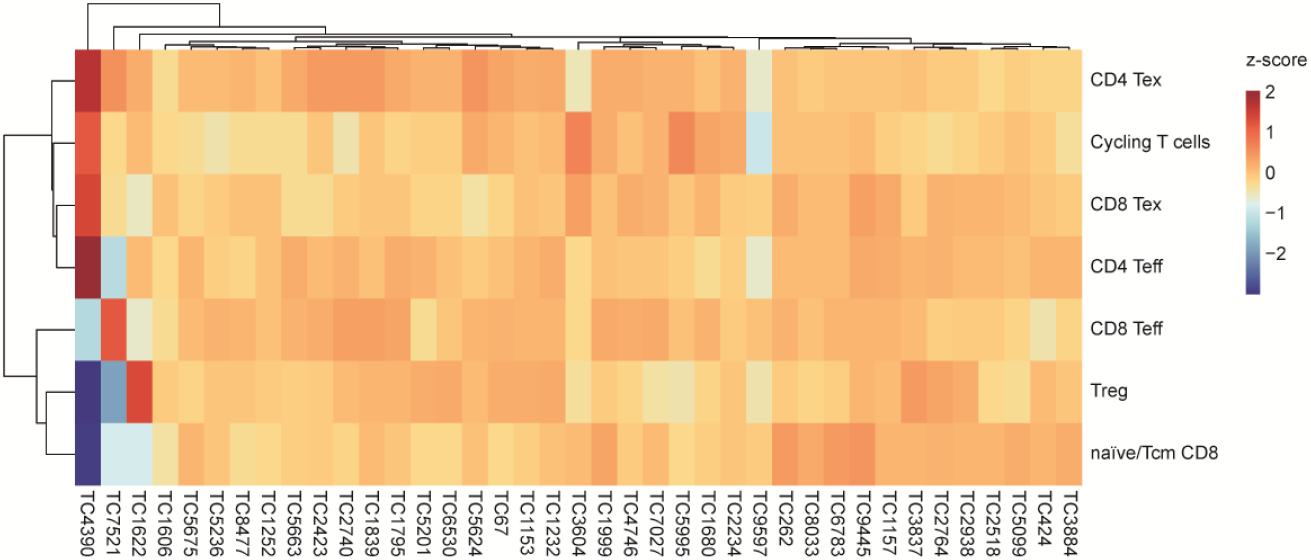
Activity of TLS-associated TCs in T cells from neoadjuvant treated EC tumors. Heatmap displaying the activity scores of 39 TCs across different the different clusters identified in the scRNAseq of tumor-infiltrating T cells in MMRd EC patients treated with anti-PD-1.

## STAR METHODS

### Human samples

The research protocol underwent evaluation by the medical ethical review board of University Medical Centre Groningen (UMCG, The Netherlands), which determined its exemption from the purview of the Medical Research Involving Human Subjects Act (WMO) (METc 2022/223). Consequently, it was reviewed and approved by the Central Ethical Review Board for non-WMO research at UMCG (202200193). The study was conducted in accordance with the protocol, the declaration of Helsinki, and the International Conference on Harmonization Guidelines for Good Clinical Practice. Eligible participants for the trial encompassed women diagnosed with endometrial cancer, spanning all grades, stages, and molecular subtypes, who had previously undergone standard surgical interventions and had their specimens archived within the pathology biobank of UMCG. In instances where molecular subtypes were not established as part of standard-of-care, analysis of MMR-expression, P53, and/or POLE mutation status was conducted.

### Method details

#### Evaluation of the presence of tertiary lymphoid structures

Tumor specimens collected as part of routine clinical care, and archived in the pathology biobank of UMCG and Leiden University Medical Center (LUMC) underwent evaluation by an expert gynecologic pathologist. Tertiary lymphoid structures (TLS) were identified using hematoxylin and eosin (H&E) stained FFPE slides, relying on histopathological and morphological characteristics. A follow-up BCL6 staining was conducted, where required, to confirm the presence of TLS. All IHC was carried out on the fully-automated Benchmark ULTRA platform (Roche, Ventana Medical Systems, Oro Valley, AZ, USA), under standardized laboratory conditions, accredited following the ISO15189 quality system.

#### Tissue processing for spatial transcriptomics

FFPE blocks were sectioned, and RNA quality was assessed. Two 10 µm slides were used for RNA isolation, according to manufacturer instructions (RNeasy FFPE kit, Qiagen). The quality of the isolated RNA was assessed using an Agilent RNA screen tape which was run on the Agilent tapestation. When the percentage of total RNA fragments > 200 nucleotides (DV200) was > 40%, RNA quality of the FFPE block was considered good enough to be used for spatial transcriptomics.

A 6.5 × 6.5 mm^2^ area from each FFPE block was selected, sectioned at thickness of 5 µM and placed on the spatial gene expression slide provided by the manufacturer. On one spatial gene expression slide, four areas (from four different FFPE blocks) could be analysed. Next, spatial transcriptomics was performed according to manufacturer instructions (10x Genomics, used protocols: CG000407, CG000408, CG000409, CG000436). The quality of the libraries was assessed on the Agilent tapestation using a High Sensitivity DNA D5000 screen tape. In each sequencing run, the four samples from one gene expression slide were pooled at an equimolar ratio. Next, 650 pM of this pool was loaded on a NextSeq2000 sequencer (with a 1% PhiX control library spiked in). A 100 bp reagent kit P2 V3 was used for the sequencing, containing 800×10e^6^ paired end reads. The read lengths were as follows: read 1 = 29 bp, read 2 = 87 bp; index 1 (i7) = 11 bp and index 2 (i5) = 11 bp.

#### Single cell isolation, sorting and 10x single-cell RNA-sequencing

Tumor tissue from endometrial cancer patients was collected during primary surgery. Tumors were minced into pieces and enzymatically digested with RPMI (Gibco) containing 1 mg/ml Collagenase type IV (Sigma) and 13 U/ml Pulmozyme (Roche), using the GentleMACS Dissociator (Miltenyi), program h_tumor_01.01. Digestion was either done overnight at room temperature or for 30 minutes at 37 °C. Digests were filtered using 70 µm cell strainers and cells were enriched using Ficoll-Paque PLUS (GE Healthcare). After washing with PBS, digests were cryopreserved in fetal bovine/calf serum supplemented with 10% dimethylsulfoxide (DMSO), and stored at liquid nitrogen until use. The day of the experiment, tumor digests were thawed in FCS and resuspended in RPMI + 10% FCS. After 1.5h incubation at 37°C/5% CO2, samples were stained and incubated with a CXCR5 PerCP-Cy5.5 antibody for an additional 0.5h at 37°C. After a total of 2h recovery at 37°C/5% CO2, samples were centrifuged and resuspended in 100 µl PBS + 2% FCS. Samples were incubated for 30-45 min on ice (in dark) with the following antibodies: CD45-BV605, CD8-APC-Cy7, CD4-PE, CD19-BV421, PD-1-APC, CD3-PE-Cy7 (Table S1). 3 µl of each antibody was used per 1×10e6 cells. Samples were washed twice with PBS + 2% FCS and filtered using a 35 µm strainer (Falcon) before sorting on a Beckman Coulter MoFlo Astrios cytometer. UltraComp Ebeads (Thermo Fisher Scientific) were used as compensation controls. Before sorting, Propidium Iodide (PI) was added to exclude dead cells. Sorting was done in 1.5 mL Low Bind DNA tubes (Eppendorf) in 150 µl PBS 0,04 % BSA. CD45+ CD3-CD19+ (B cells), CD45+ CD3+ CD19-CD8+ (CD8 cells) and CD4+ (CD4 cells) or CD45+ CD3-CD19- (other CD45 cells). Sorted cells were centrifuged and resuspended at a concentration of 727 – 2121 cells/µl (total 16.5 µl) in PBS 0,04% BSA, depending on the total number of sorted cells and the sample. For samples where the targeted number of B cells could not be sorted, the sample was pooled with cells from the ‘non-B/non-T CD45’ subset, and for samples where the target number of CD4 or CD8 T cells could be reached, both CD4 and CD8 were pooled together at equal ratios. As such, the fraction of single cells sequenced is not a relative reflection of the immune cell composition, but an enriched pool to capture as much immune cell diversity as possible. Subsequent steps were performed according to manufacturer’s protocol (10x genomics): ‘chromium next GEM single cell 5’ reagents kit v2 (dual index)’. For the cDNA amplification step, 13 cycles were done. Although not analyzed and reported in the current work, VDJ amplification was performed in parallel, and either the B cell kit or T cell kit was used, depending on the sorted population (B cells or T cells). For each sample, the gene expression (GEX) library was constructed. All cDNA quality control and quantification steps were done with the Qubit 4 Fluorometer (Thermo Fisher) and the Agilent 4200 Tapestation (with D5000 and high sensitivity D5000 screen tapes). For the sequencing, V(D)J and GEX libraries were pooled at a 1:4 molar ratio, with 2 GEX libraries and 2 V(D)J libraries in one pool. 1.5 pM of this superpool was sequenced on a NextSeq 500 (Illumina inc., San Diego, CA, USA). A 150 bp NextSeq high-output reagent kit was used for the sequencing (paired end).

#### Sequencing read alignments, quality control, cell clustering, and annotation

Reads from spatial transcriptomics and single cells isolated using 10x chromium were demultiplexed and aligned to the GRCh38.p12 human reference genome (from 10x Genomics) using Cell Ranger (version 6.0.1; 10x Genomics)^70^. Cell Ranger outputs were loaded into R (Seurat (V4) package)^71^.

Spatial Transcriptomic Seurat objects from different patients were integrated using sequentially the ‘SCTransform’ (assay = ‘Spatial’, vst.flavor = ‘v2’, and method = ‘glmGamPoi’), ‘SelectIntegrationFeatures’ (n=3000), ‘PrepSCTIntegration’,’ ‘RunPCA’, FindIntegrationAnchors’ (utilizing 50 dimensions, rpca as reduction, SCT as normalization method, and k.anchor set to 20), and ‘IntegrateData’ (utilizing 50 dimensions with SCT as normalization method) commands. TCR and BCR genes were removed from the variable features of the merged Seurat object. Principle components of the merged Seurat object were calculated using the ‘RunPCA’ command. Global clusters of similar spatial regions (spots) were detected using the Louvain method for community detection to construct the shared nearest neighbour map and an empirically set resolution using the ‘FindNeighbours’ (utilizing 50 dimensions) and ‘FindClusters’ (resolution 1) commands. Spatial Transcriptomic regions were annotated by expert pathologic review.

Single cells with fewer than 200 genes, more than 4000 genes, or >10% mitochondrial counts were excluded from further analysis. Seurat objects from different patients were subsequently integrated using sequentially the ‘NormalizeData’, ‘FindVariableFeatures’, ‘ScaleData’, ‘RunPCA’, ‘FindIntegrationAnchors’ (utilizing 50 dimensions with rpca as reduction), and ‘IntegrateData’ commands (utilizing 50 dimensions). Expression values of the merged Seurat object were converted to z-scores using the ‘ScaleData’ command. Principle components were calculated using the ‘RunPCA’ command. Global clusters of similar cells were detected using the Louvain method for community detection to construct the shared nearest neighbour map and an empirically set resolution using the ‘FindNeighbours’ (utilizing 50 dimensions) and ‘FindClusters’ (resolution 0.1) commands. Cells were annotated using previously reported canonical markers (Figure S2). Heterotypic clusters expressing more than 1 mutually exclusive canonical marker (n=3) were excluded.

For granular analysis of endothelial cells, the endothelial-annotated cluster was subset and principle components recalculated using the ‘RunPCA’ command. Global clusters of similar endothelial cells were detected using the Louvain method for community detection to construct the shared nearest neighbour map and an empirically set resolution using the ‘FindNeighbours’ (utilizing 50 dimensions) and ‘FindClusters’ (resolution 1) commands. Cells were annotated using previously reported canonical markers.

#### Celltype inference, radial distance, region segmentation, cell cycle score, gini-TCR and transcription factor activity

Cell Ranger outputs were loaded into R (Semla (version 1.1.6)) using sequentially the ‘ReadVisiumData’ and ‘LoadImages’ (with image height set to 1904) commands. Cell identities were carried over from the merged Spatial Transcriptomics and single cell Seurat objects using the spot/cell barcodes. Celltype inference was performed using the ‘RunNNLS’ command. Radial distances (in micron) from the TLS cluster calculated using the ‘RadialDistance’ command. Individual TLS per selected using sequentially the ‘RegionNeighbors’, and ‘DisconnectRegions’ commands, removing single TLS regions that were characterized by only a single spatial spot (‘singleton’ output from ‘DisconnectRegions’). Cell cycle scores were calculated using the ‘CellCycleScoring’ command from the Seurat package. Gini-TCR scores were calculated using the ‘ineq’ package.

Activity of transcription factors was inferred using the decoupleR (v2.4.0) and DoRothEA (v1.10.0) packages with the “run_wmean” command (times set to 100 and minsize to 5). Only networks with the highest confidence levels were used for the analysis (level ‘A’). Activities were scaled using the Seurat “ScaleData” command, mean activities calculated by cluster, and the transcription factors most variable across clusters selected (maximum of 50). Top transcription factor activities were visualized using the “pheatmap” command from pheatmap package (clustered using the ward.D2 method). A DC help signature score was calculated using the “AddModuleScore_UCell” command from the UCell package (v2.2.0) and a previously reported CD4 help signature gene list.

Versions:

viridis_0.6.3 for the spatial gene expression plots

ggplot2_3.4.2 and ggpubr_0.6.0 for the rest of the figures

Seurat_4.3.0

semla_1.0.0

#### Analysis of tumor-draining lymph nodes

Spatial transcriptomic reads from tumor-draining lymph nodes and tonsil were processed as described above for tumor tissue. In brief, reads were demultiplexed, aligned to the human reference genome, loaded into R and integrated into a single Seurat object. TCR and BCR genes were removed from the variable features, principle components calculated, and global clusters of similar spatial regions detected using the Louvain method for community detection to construct the shared nearest neighbour map (resolution 1 as used for tumors). Spatial Transcriptomic regions were annotated using canonical markers combined with pathology review. G2M and S scores were calculated for GC spots as described above. TLS and TDLN-GC clusters from different patients were integrated using sequentially the ‘SCTransform’ (assay = ‘Spatial’, vst.flavor = ‘v2’, and method = ‘glmGamPoi’), ‘SelectIntegrationFeatures’ (n=1000), ‘PrepSCTIntegration’,’ ‘RunPCA’, FindIntegrationAnchors’ (utilizing 50 dimensions, rpca as reduction, SCT as normalization method, and k.anchor set to 20), and ‘IntegrateData’ (utilizing 50 dimensions with SCT as normalization method) commands. TCR and BCR genes were removed from the variable features of the merged Seurat object. BCL6 and CD21 staining was carried out on the fully-automated Benchmark ULTRA platform (Roche, Ventana Medical Systems, Oro Valley, AZ, USA), under standardized laboratory conditions, accredited following the ISO15189 quality system.

#### Statistical analysis of spatial and single cell transcriptomics

Differentially expressed genes between clusters were calculated using the ‘FindAllMarkers’ command and visualized using the ‘DoHeatmap’ command, both from the Seurat package (version as above). Pairwise differences between multiple groups were calculated using the ‘Stats’ package with the ‘pairwise.wilcox.test’ command with benjamini hochberg correction for multiple testing.

#### Immunohistochemistry and immunofluorescence

IHC stainings for IgA, IgG, BCL6, and CD21 staining were carried out on the fully-automated Benchmark ULTRA platform (Roche, Ventana Medical Systems, Oro Valley, AZ, USA), under standardized laboratory conditions, accredited following the ISO15189 quality system. In short, paraffin tissue sections (3 µm) were incubated with antibodies against IgA (1:400, Cell Marque, polyclonal), IgG (R.T.U., Ventana, polyclonal), BCL6 (R.T.U., Ventana, CI191E/A8), CD21 (R.T.U., Ventana, 2G9). Each slide contained a suitable tissue section, serving as external control. For immunofluorescence staining, tissue slides were deparaffinized and rehydrated, followed by heat-induced epitope retrieval (10 mM citrate buffer pH 6) and blocking of endogenous peroxidase (0.3% H2O2 blocking buffer). Tissue slides were incubated overnight with rabbit anti-human FCRL3 antibody (1:250, HPA048022, Sigma-Aldrich) followed by incubation with Envision+HRP anti-rabbit (K4003, Agilent) and subsequently TSA Cyanine 5 detection solution (NEL705A001KT, Perkin Elmer) according to manufacturer’s instructions. HRP labels were destroyed by incubating tissue slides in 0.01 M HCl for 10 min at RT. Slides were then incubated overnight with mouse anti-human CD20 antibody (1:200, clone L26, M075501, Agilent) followed by incubation with Envision+HRP anti-mouse (K4001, Agilent) and subsequently TSA Plus Cyanine 3 detection solution (NEL753001KT, Perkin Elmer) according to manufacturer’s instructions. HRP labels were again destroyed by incubating tissue slides in 0.01 M HCl for 10 min at RT. Slides were then incubated overnight with rabbit anti-human HRP-conjugated CD21 antibody (1:300, clone EP3093, ab202354, Abcam) followed by incubation with TSA Plus Fluorescein detection solution (NEL753001KT, Perkin Elmer) according to manufacturer’s instructions. Tissue slides were embedded in ProLong Diamond Antifade Mountant with DAPI (P36962, ThermoFisher Scientific). Images were made with a ZEISS Axio Imager 2 microscope.

#### TCGA data acquisition

From TCGA, we obtained the pre-processed and normalized level 3 RNA-seq (version 2) data for 27 cancer datasets available at the Broad GDAC Firehose portal (downloaded January 2017 https://gdac.broadinstitute.org/). For each sample, we downloaded RNA-Seq with Expectation Maximization (RSEM) gene normalized data (identifier: illuminahiseq_rnaseqv2 RSEM_genes_normalized according to NCI Genomic Data Commons (GDC) User’s Guide). The complete workflow of the transcriptomic analysis can be found in figure 4A.

#### Consensus-independent component analysis (c-ICA)

The bulk transcriptome input data underwent a whitening transformation to optimize the subsequent analysis. Following this pre-processing step, we employed the FastICA algorithm to execute the independent component analysis, facilitating the derivation of estimated sources (ESs)^72,73^. We determined the appropriate number of ESs to extract based on the number of principal components that captured 100% of the total variance within the dataset. To rigorously evaluate the stability of the ESs, we conducted 25 separate ICA runs, each with uniquely randomized initial weight factors. ESs exhibiting an absolute Pearson correlation greater than 0.9 were grouped together. From these groupings, we derived consensus ESs, referred to as transcriptional components (TCs), by selecting the ES with the least correlation to ESs outside its group. Only groups with at least two member ESs were considered in this analysis. This approach is based on the hypothesis that any ES appearing in at least two runs is likely to be a non-random ES, as the probability of a random signal appearing in two different runs is minimal. The gene weights within each TC provide insights into the strength and directionality by which an underlying latent biological process influences the related gene expression levels. Subsequently, these TCs are used to generate a consensus mixing matrix (MM). The coefficients within this MM depict the activity scores of the TCs across the samples.

#### Group-wise comparison

To discern the relationship between TC activity and TLS presence, a Mann-Whitney U test was conducted on a select group of patients with available information. We implemented a permutation-based framework encompassing 10,000 permutations to mitigate the risk of false discoveries. We established the acceptable false discovery rate (FDR) at 5%, maintaining an 80% confidence level, applicable for both the Mann-Whitney U test.

#### Associating the identified transcriptional components with biological processes

To determine the biological processes linked to the transcriptional components (TCs), we employed a comprehensive approach that included: i) Transcriptional Adaptation to Copy Number Alterations (TACNA) Profiling: This step focused on identifying TCs that reflect the downstream effects of copy number alterations (CNAs) on gene expression levels^73^. ii) Gene Set Enrichment Analysis (GSEA): For each TC, we conducted GSEA using 16 gene set collections from The Human Phenotype Ontology (The Monarch Initiative), the Mammalian Phenotypes (Mouse Genome Database), and the Molecular Signatures Database (MsigDB)^42,43^. Iii) Co-Functionality Networks Formation: We constructed co-functionality networks for the top and bottom genes of each TC using the GenetICA methodology^74^, available at https://www.genetica-network.com. For clusters containing five or more genes, we quantified the enrichment of the predicted functionality. This enrichment served as the basis for identifying the biological processes associated with each TC.

#### Determining activity of transcriptional components in spatial and single cell transcriptomes

In-house spatial resolved transcriptomic profiles and single cell transcriptomic profiles from endometrial cancer samples were obtained for analysis. Activity for each TC across every location within the spatial samples and single cell profiles was ascertained through the cross-study projection methodology referred to in the previous study^71^. To discern the markedly active areas within the spatial or single cell samples for each TC, we incorporated a permutation-driven approach. We derived a null-distribution of activities for each TC-profile pairing by performing 3,000 permutations and subsequent projection. For every null distribution, if the null distribution does not adhere to a Gaussian distribution, as determined by the Anderson-Darling test, we apply the Johnson transformation to convert the null distribution into a Gaussian distribution. This transformation employs one of three optimal families of distributions (S, SU, SL) and finds the parameters that transform the null distribution as much as possible to a Gaussian distribution. Apply this transformation also to the corresponding original activity score. We then fit a symmetrical, generalized Gaussian distribution to the null distribution. Next, we extract the p-value of the transformed activity score and convert it to a Z-score to obtain the final normalized activity score.

