## Supplementary methods for "The landscape of tertiary lymphoid structures in endometrial cancer revealed through harmonized multi-level transcriptomics"

**Consensus Independent Component Analysis (c-ICA)**

Independent Component Analysis (ICA) is a statistical method to decompose mixed multivariate signals into their constituent source signals. It operates under the assumption that these source signals, termed independent components (ICs), are statistically independent and follow non-Gaussian distributions. This study applies ICA to bulk transcriptional profiles (each containing measurements for $p$ genes), considered mixed multivariate signals, to isolate unique source transcriptional patterns (i.e., ICs). These patterns are indicative of distinct underlying biological processes. Each IC comprises $p$ gene weights, representing the direction and magnitude of the effect of the underlying biological process on a gene's expression level.

Our c-ICA approach involves four steps: A) Whitening the bulk transcriptional profiles; B) Applying Independent Component Analysis on the whitened transcriptional profiles; C) Consensus approach; D) Normalizing the consensus mixing matrix. The steps are described in detail below.

*A) Whitening the bulk transcriptional profiles*

Whitening the bulk transcriptional profiles is an essential preprocessing step before applying ICA, where we transform the matrix $\boldsymbol{X}$ containing the bulk transcriptional profiles in rows and measurements for $p$ genes in the columns into a new matrix $\boldsymbol{X}_{whitened}$ with whitened transcriptional profiles. This transformation gives the whitened transcriptional profiles desirable statistical properties:

1) Unit variance: This standardizes the scale of each whitened transcriptional profile, ensuring that each has a variance of 1. As a result, all whitened transcriptional profiles are on a comparable scale, which is critical for the subsequent analysis because it prevents any single whitened transcriptional profile from dominating the results due to differences in the magnitude of gene expression.
2) Zero covariance: By transforming the bulk transcriptional profiles so that the covariance between any pair is zero, we ensure they are uncorrelated. In the covariance matrix of $\boldsymbol{X}_{whitened}$ this is reflected by having diagonal elements representing variances (now one due to unit variance scaling) and off-diagonal elements (representing covariances between different profiles) being zero.
3) Orthogonality: With zero covariance, the whitened transcriptional profiles also achieve orthogonality, meaning they are perpendicular to each other in the multidimensional gene expression space. This orthogonality provides a robust basis for separating the whitened transcriptional profiles into independent components during the ICA process.

The process of whitening is critical when performed before applying ICA to bulk transcriptional profiles for two main reasons:

1) Enhancing the efficiency of ICA: As described above, whitening transforms the matrix of bulk transcriptional profiles, denoted as $\boldsymbol{X}$, into a new matrix $\boldsymbol{X}_{whitened}$ with desirable statistical properties. Such a transformation simplifies the optimization landscape for ICA, making it smoother and more tractable. Consequently, this increases the efficiency of the ICA algorithm, leading to faster convergence.
2) Reducing dimensionality and mitigating noise: Beyond expediting convergence, whitening can play a role in reducing the dimensionality of the data and mitigating noise. By transforming the bulk transcriptional profiles into an orthogonal and uncorrelated space, it becomes feasible to pinpoint and disregard whitened transcriptional profiles that contribute minimal variance, which often represent noise or less informative transcriptional patterns in the data. Discarding these whitened transcriptional profiles reduces computational demand and enhances the ability of ICA to discern transcriptional patterns that reflect distinct biological processes.

Whitening the matrix $\boldsymbol{X}$ containing the bulk transcriptional profiles involves the following steps:

1) Center each profile: To center each bulk transcriptional profile in matrix $\boldsymbol{X}$, subtract the mean of its respective row. The centered matrix $\tilde{\boldsymbol{X}}$ is obtained by the operation $\tilde{\boldsymbol{X}}=\boldsymbol{X}-{\mu1}^{T}$, where $\mu$ is the vector of row means, and $1^{T}$ is a row vector of ones with a length matching the number of columns in $\boldsymbol{X}$. This process adjusts each value in a row by the row's mean, centering the bulk transcriptional profiles.
2) Calculate the covariance matrix: Compute the covariance matrix $\boldsymbol{C}$ of the matrix$\tilde{\boldsymbol{X}}$. The covariance is computed as $\boldsymbol{C}= \frac{1}{N-1}\tilde{\boldsymbol{X}}\boldsymbol{X}^{T}$, where $\boldsymbol{X}^{T}$ is the transpose of$\tilde{\boldsymbol{X}}$, and $N$ is the number of genes (columns of $\tilde{\boldsymbol{X}}$).
3) Eigenvalue decomposition: Perform an eigenvalue decomposition on the covariance matrix $\boldsymbol{C}$. Since $\boldsymbol{C}$ is a positive semi-definite matrix, it can be decomposed $\boldsymbol{C}=\boldsymbol{V}\boldsymbol{\Lambda}\boldsymbol{V}^{T}$, where $\boldsymbol{V}$ is a matrix of eigenvectors and $\boldsymbol{\Lambda}$ is a diagonal matrix with eigenvalues on the diagonal. The eigenvectors correspond to the principal axes of the data distribution, and the eigenvalues correspond to the variance explained by each axis.
4) Form the whitening transform: Create the whitening transform matrix $\boldsymbol{W}$ using the eigenvalues and eigenvectors. Each eigenvalue $\lambda_{i}$ is used to form a scaling factor $\lambda_{i}^{-1/2}$ (assuming $\lambda_{i}$ is non-zero). The whitening matrix $\boldsymbol{W}$ is then constructed as $\boldsymbol{W}=\boldsymbol{V}\boldsymbol{\Lambda}^{-1/2}\boldsymbol{V}^{T}$, where $\boldsymbol{\Lambda}^{-1/2}$is the diagonal matrix of the inverse square roots of the eigenvalues.
5) Apply the whitening transform: Multiply the matrix $\tilde{\boldsymbol{X}}$ by the whitening matrix $\boldsymbol{W}$ to obtain the whitened matrix $\boldsymbol{X}_{whitened}=\tilde{\boldsymbol{X}}\boldsymbol{W}$. The result is that the covariance matrix of $\boldsymbol{X}_{whitened}$ is the identity matrix $\boldsymbol{I}$, meaning that the rows of $\boldsymbol{X}_{whitened}$ are linearly uncorrelated/orthogonal, and each has unit variance.
6) Dimensionality and noise reduction (optional): You can choose a subset of the eigenvectors—for example, corresponding to the largest eigenvalues—before constructing $\boldsymbol{W}$. This effectively reduces the number of dimensions in the whitened data, as you're only keeping the eigenvectors with the most variance, which typically carry the most information. This automatically discards the eigenvectors that contribute the least variance. These eigenvectors often represent noise or less informative features of the data.

In the whitening process, there's an important setting known as the *Cumulative Explained Variance Threshold*. This setting allows we to specify the amount of the original data's variance we want to preserve in the whitened data. The cumulative explained variance is the total variance accounted for by the selected eigenvectors. Choosing a cumulative explained variance threshold—like 90%—determines that we want to keep enough eigenvectors to capture 90% of the original variance. In practice, we would add up the eigenvalues from largest to smallest until the sum is equal to or just exceeds 90% of the sum of all eigenvalues. The minimum number of eigenvalues we add up to meet this threshold is the number we keep. The remaining eigenvectors, which correspond to smaller eigenvalues and thus less variance, are discarded. As described above, this approach ensures that the whitened matrix $\boldsymbol{X}_{whitened}$ maintains the majority of the informative variance from the original matrix while reducing dimensionality and potentially removing noise. This matrix $\boldsymbol{X}_{whitened}$ with whitened transcriptional profiles serves as input to ICA.

*B) Applying Independent Component Analysis on the whitened transcriptional profiles*

ICA is performed with the fastICA algorithm. FastICA is an iterative, fixed-point algorithm designed for the task of ICA. It reaches a solution through an iterative process that converges to a fixed point where the algorithm's output becomes stable and does not change significantly with further iterations. A detailed description of the fastICA algorithm has been provided by Hyvarinen et al. ^1^ In brief, the algorithm operates under the assumption that the observed bulk transcriptional profiles are linear mixtures of statistically independent, non-Gaussian source transcriptional patterns. The primary objective is to estimate the unmixing matrix that recovers these statistically independent source transcriptional patterns (i.e., ICs) when applied to the observed bulk transcriptional profiles.

Four key parameters govern FastICA:

1) *Mu* ($\mu$): This parameter is often referred to as the learning rate or step size in the context of iterative optimization algorithms. In FastICA, *Mu* controls the magnitude of the updates made to the unmixing matrix in each iteration. An adequately chosen *Mu* is crucial for the algorithm's convergence; too large a *Mu* may cause the algorithm to oscillate or diverge, while too small a *Mu* can lead to very slow convergence.
2) *Contrast Function* ($g$): The contrast function is a key component in ICA algorithms that measures non-Gaussianity. The contrast function maximizes the non-Gaussianity of the estimated source transcriptional patterns (i.e., ICs), assuming that the true source transcriptional patterns are statistically independent and have non-Gaussian distributions. Different choices of contrast functions can lead to different fastICA solutions, and the choice of function may depend on the nature of the true source transcriptional patterns. Common examples include hyperbolic tangent, kurtosis, raise to power 3, and skewness. The contrast function directly influences the update rule applied to the unmixing matrix during optimization.
3) *Epsilon* ($\varepsilon$): This parameter is a small positive value that serves as a convergence criterion. Specifically, *Epsilon* is used to determine when the change in the non-Gaussianity metric between consecutive iterations is small enough to consider the algorithm to have converged. In other words, when the algorithm's updates lead to changes smaller than *Epsilon*, the algorithm stops iterating, assuming that further iterations will not lead to significant improvements. This parameter helps to prevent infinite loops in cases where exact convergence to a threshold is not achievable.
4) *Maximum Number of Iterations* (${max}_{iter})$: This parameter sets an upper limit on the number of iterations the algorithm will perform. This safeguards against non-convergence or extremely slow convergence, ensuring that the algorithm terminates after a reasonable time even if the *Epsilon* convergence criterion has not been met. The choice of this value may depend on the application context and the computational resources available; a higher value allows for more thorough convergence at the cost of increased computational time.

FastICA employs a random initialization for the unmixing matrix and refines the unmixing matrix through an iterative process. Each iteration consists of two main steps: calculating the unmixing matrix update based on a chosen non-linear contrast function and the subsequent orthogonalization of the updated unmixing matrix to ensure that the estimated source transcriptional patterns remain statistically independent. The iterative process continues until a convergence criterion is met, defined as the change in the unmixing matrix between successive iterations falling below our predetermined threshold *Epsilon*. This criterion ensures that the algorithm terminates once the estimated source transcriptional patterns are sufficiently statistically independent and the unmixing matrix stabilizes. Upon convergence, the FastICA algorithm has decomposed the observed bulk transcriptional profiles into their constituent statistically independent source transcriptional patterns (i.e., ICs).

In formulas:

Given $\boldsymbol{X}_{whitened}$, which is the matrix containing the whitened transcriptional profiles, estimate the unmixing matrix $\boldsymbol{W}$ such that: $\boldsymbol{S}=\boldsymbol{W}\boldsymbol{X}_{whitened}$. Here $\boldsymbol{S}$ is a matrix containing the independent source transcriptional patterns (i.e., ICs), which are assumed to be non-Gaussian and statistically independent. The FastICA algorithm proceeds iteratively as follows:

1) Initialization: Choose an initial random unmixing matrix $\boldsymbol{W}$.
2) Iteration:
   a. For each component $i$, update $\boldsymbol{W}$ using the fixed-point iteration scheme:

$$w_{i}^{new}=w_{i}+\mu(\mathbb{E}\left[ \boldsymbol{X}_{whitened}\cdot g\left( w_{i}^{T}\boldsymbol{X}_{whitened} \right) \right]- \mathbb{E}\left[ g^{'}\left( w_{i}^{T}\boldsymbol{X}_{whitened} \right) \right]\cdot w_{i})$$

Where:

- $w_{i}$ is the $i^{th}$ row of $\boldsymbol{W}$.
- $g(\cdot)$ is the non-linear contrast function and $g^{'}\left( \cdot\right)$ its derivate.
- $\mathbb{E}\left[ \cdot\right]$ denotes the expected value (mean).

3) Normalization: Normalize $w_{i}^{new}$: $w_{i}^{new}=\frac{w_{i}^{new}}{\parallel w_{i}^{new}\parallel}$.
4) Orthogonalization: Ensure that the components remain linearly uncorrelated by orthogonalizing the vectors: $\boldsymbol{W}^{new}=orthogonalize(\boldsymbol{W}^{new})$.
5) Convergence and itereration check: Check if the change in $\boldsymbol{W}$ is below the threshold $\varepsilon$. If it is, assume convergence: $\parallel\boldsymbol{W}^{new}-\boldsymbol{W}\parallel<\varepsilon$. If the convergence criterion is not met, set $\boldsymbol{W}= \boldsymbol{W}^{new}$and repeat the iteration process if ${max}_{iter}$ is not reached yet.
6) End: Once convergence is reached, the final unmixing matrix $\boldsymbol{W}$ can be used to compute the statistically independent components $\boldsymbol{S}=\boldsymbol{W}\boldsymbol{X}_{whitened}$.

*C) Consensus approach*

The fastICA algorithm employs an optimization technique to decompose whitened transcriptional profiles of mixed signals $\boldsymbol{X}_{whitened}$ into statistically independent source transcriptional patterns (i.e., ICs) $\boldsymbol{S}$. However, this optimization process can converge to local optima, particularly in high-dimensional or noisy data spaces, leading to solutions that are not globally optimal. These local solutions may vary with different random initializations of the unmixing matrix $\boldsymbol{W}$, resulting in different sets of ICs $\boldsymbol{S}_{run}$ upon each run of the algorithm.

A consensus approach addresses this challenge by aggregating the results from multiple runs of the fastICA algorithm, each with a different random initialized unmixing matrix $\boldsymbol{W}$ and identifying the ICs extracted across multiple runs. This approach aims to filter out ICs likely to be noise artifacts specific to a particular run or from convergence to a suboptimal local solution. The consensus approach enhances reliability by focusing on the stable and robust ICs that emerge across multiple runs.

Three parameters govern the consensus approach:

1) *Consensus Runs*: This parameter dictates how often the FastICA algorithm will be executed. In each run, the unmixing matrix $\boldsymbol{W}$ is randomly initialized with a different random seed, affecting the optimization process's starting conditions. The significance of this parameter lies in its ability to capture a wide range of potential solutions, which helps determine the consistency of the ICs identified across runs. A higher number of runs increases the chances of capturing stable and robust ICs, allowing for a more comprehensive sampling of the solution space. However, this also increases computational load.
2) *Consensus threshold*: This parameter specifies the minimum Pearson correlation coefficient ICs must exceed to be considered equivalent across different runs. A higher correlation threshold requires a stronger linear relationship between ICs to be regarded as the same. This could lead to identifying only the most consistent and robust ICs across runs. However, this might also dismiss less correlated but potentially relevant ICs, thereby reducing the sensitivity of the analysis.
3) *Credibility index*: This parameter serves as a threshold to determine whether a consensus-independent component (c-ICs) is robust enough to be considered reliable. It measures how consistently an IC appears across different runs. If a IC is identified in a proportion of runs that exceeds the credibility index, it is deemed robust. The credibility index filters out ICs that are less likely to be true signals and more likely to be noise or artifacts, as these would not consistently appear across multiple runs. The credibility index's choice impacts the filtering process's strictness: setting it too high might exclude genuine ICs. At the same time, a too low threshold might include spurious ones.

The consensus approach involves several steps:

1) Consolidation of ICs: Initially, the ICs $\boldsymbol{S}_{run}$ derived from each run are combined to form a matrix $\boldsymbol{S}_{combined}$.
2) Computation of pair-wise correlations: Begin by centering each IC $S$ in $\boldsymbol{S}_{combined}$ subtracting its mean and normalizing it to its L2-norm resulting in the matrix ${\tilde{\boldsymbol{S}}}_{combined}$. Then, calculate the product of the matrix ${\tilde{\boldsymbol{S}}}_{combined}$ with its transpose: ${\tilde{\boldsymbol{S}}}_{combined}{{(\tilde{\boldsymbol{S}}}_{combined})}^{T}$. The resulting matrix $\boldsymbol{R}$ will have diagonal elements that signify the self-correlation of each IC $S$, and off-diagonal elements that indicate the Pearson correlation coefficients between different ICs. The absolute value of these coefficients is then used to evaluate the similarity between the ICs.
3) Establishment of sets of linked ICs: For each IC $S$ in the matrix $\boldsymbol{S}_{combined}$, the correlation matrix $\boldsymbol{R}$ is utilized to find ICs correlated with $S$ above the *Consensus Threshold*. Those that exceed this threshold are deemed equivalent. ICs $S$ and these equivalent ICs constitute a set of linked ICs $L_{set}$. Subsequently, IC $S$ and its corresponding set $L_{set}$ are paired as key-value and added to $\boldsymbol{L}_{map}$, where $S$ serves as the key and $L_{set}$ as the value.
4) Organize ICs into consensus clusters: An iterative process $I$ for forming consensus clusters is started by processing $\boldsymbol{L}_{map}$:
   - Take the largest set of linked ICs $L_{{set}_{largest}}$ from $\boldsymbol{L}_{map}$ and use its ICs to create a new consensus cluster.
   - Then, a recursive process $P$ starts to update $\boldsymbol{L}_{map}$ that takes a set of linked ICs $L_{set}$ as parameter $Lparam$. This recursive process is started with $L_{{set}_{largest}}$ as parameter $Lparam$. In this recursive process $P$ the following steps are performed:
     1. Eliminate any key-value pairs from $\boldsymbol{L}_{map}$ where the key is identical to any IC $S$ found in $Lparam$.
     2. From the $\boldsymbol{L}_{map}$, for each pair that remains, update the pair’s set of linked ICs $L_{set}$, by excluding any IC $S$ that is included in $Lparam$.
     3. For each IC $S$ within $Lparam$, we search $\boldsymbol{L}_{map}$ for a corresponding key that matches $S$. If no matching key is found, the current recursive process halts. Otherwise, if a matching pair is identified, we start another recursive process $P$, this time using the linked IC set $L_{set}$ from the matching pair as the new $Lparam$.
   - The iterative process $I$ is stopped when there are no key-value pairs left in $\boldsymbol{L}_{map}$*.* Otherwise, another iteration of $I$ is started with the current $L_{{set}_{largest}}$ from $\boldsymbol{L}_{map}$.
5) Filtering consensus clusters based on credibility index: We iterate over the consensus clusters, computing the credibility index for each. This index represents the ratio of the number of ICs within the clusters to the total number of runs (*consensus runs*). Clusters with a credibility index meeting or exceeding the *consensus threshold* are considered valid; the rest are removed. Suppose the total number of valid consensus clusters surpasses the number of whitened variables. In that case, we only retain the $N$clusters with the largest credibility index, where $N$ equals the number of whitened variables.
6) Determining c-ICs: For all ICs within a consensus cluster that persisted after the credibility index-based filtering, we compute a correlation matrix as described in *2) Computation of pair-wise correlations*. Then, for each consensus cluster, we identify the most representative IC (the “winner”) by identifying the IC with the lowest average correlation to those outside its cluster. We refer to this winner as the c-IC.
7) Flipping based on skewness: We determine the skewness of the gene weights for each IC. If the skewness is negative, we flip all signs of the gene weights.

*C) Calculating the consensus mixing matrix*

As described above, ICA operates under the assumption that the observed bulk transcriptional profiles can be understood as linear mixtures of source transcriptional patterns. These mixtures are composed of statistically independent, non-Gaussian source transcriptional patterns, which are encapsulated by what we refer to as c-ICs. Consequently, each bulk transcriptional profile observed can be considered a composite, where the c-ICs contribute with varying weights. These weights are systematically arranged in what is known as a consensus mixing matrix, denoted by $\boldsymbol{MM}_{consensus}$. In the $\boldsymbol{MM}_{consensus}$, each weight ${\boldsymbol{MM}_{consensus}}_{(S_{consensus}, X)}$ represents the ‘activity’ of a c-IC $S_{consensus}$ in a bulk transcriptional profile $X$. In practical terms, to derive the $\boldsymbol{MM}_{consensus}$ one must take the matrix $\boldsymbol{X}$, which contains the bulk transcriptional profiles, and perform a matrix multiplication with the pseudo-inverse of the c-ICs $\boldsymbol{S}_{consensus}$. This operation is mathematically represented as $\boldsymbol{MM}_{consensus}$= ${\boldsymbol{X}\left( \boldsymbol{S}_{consensus} \right)}^{-1}$.

*C) Normalizing the consensus mixing matrix*

In the analysis of bulk transcriptional profiles, we observe considerable variation in gene expression levels. This variation can often be attributed to technical factors such as platform inconsistencies or batch effects, rather than underlying biological differences. Additionally, within each c-IC, the number of genes carrying heigh weights varies, which could introduce bias into the calculated consensus mixing matrix, $\boldsymbol{MM}_{consensus}$. Such biases hamper direct comparisons within the $\boldsymbol{MM}_{consensus}$, whether comparing values from different profiles for a single component or across multiple components for a single profile. To address these challenges and facilitate more accurate comparisons, we have implemented a normalization methodology for the $\boldsymbol{MM}_{consensus}$:

1) Randomly permute the rows (containing genes) within the pseudo-inverse of the consensus-independent component to get $\left( \boldsymbol{S}_{{consensus}_{permuted}} \right)^{-1}$and compute ${\boldsymbol{MM}_{consensus}}_{permuted}$=${\boldsymbol{X}\left( \boldsymbol{S}_{{consensus}_{permuted}} \right)}^{-1}$. For each weight ${{\boldsymbol{MM}_{consensus}}_{permuted}}_{(S_{consensus}, X)}$, add the weight to the null distribution ${null distribution}_{(S_{consensus}, X)}$. Repeat this process for a specified number of permutations.
2) For every null distribution ${null distribution}_{(S_{consensus}, X)}$, if the null distribution does not adhere to a Gaussian distribution, as determined by the Anderson-Darling test, we apply the Johnson transformation to convert the null distribution into a Gaussian distribution ${{null distribution}_{Jtransformed}}_{(S_{consensus}, X)}$. This transformation employs one of three optimal families of distributions (S, SU, SL) and finds the parameters that transform the null distribution as much as possible to a Gaussian distribution. Apply this transformation also to the corresponding ${\boldsymbol{MM}_{consensus}}_{(S_{consensus}, X)}$ to get ${\boldsymbol{MM}_{{consensus}_{Jtransformed}}}_{(S_{consensus}, X)}$.
3) We then fit a symmetrical, generalized Gaussian distribution to ${{null distribution}_{Jtransformed}}_{(S_{consensus}, X)}$. Next, we extract the p-value of the ${\boldsymbol{MM}_{{consensus}_{transformed}}}_{(S_{consensus}, X)}$and convert it to a Z-score to obtain the final normalized mixing matrix weight ${\boldsymbol{MM}_{normalized}}_{(S_{consensus}, X)}$.
